# Genome-scale sequencing and analysis of human, wolf and bison DNA from 25,000 year-old sediment

**DOI:** 10.1101/2021.01.08.425895

**Authors:** Pere Gelabert, Susanna Sawyer, Anders Bergström, Thomas C. Collin, Tengiz Meshveliani, Anna Belfer-Cohen, David Lordkipanidze, Nino Jakeli, Zinovi Matskevich, Guy Bar-Oz, Daniel M. Fernandes, Olivia Cheronet, Kadir T. Özdoğan, Victoria Oberreiter, Robin N. M. Feeney, Mareike C. Stahlschmidt, Pontus Skoglund, Ron Pinhasi

**Affiliations:** Department of Anthropology, University of Vienna, Vienna, Austria; Ancient Genomics Laboratory, Francis Crick Institute, NW1 1AT London, United Kingdom; School of Medicine, University College Dublin, Dublin 4, Ireland; Georgian National Museum, Institute of Paleoanthropology and Paleobiology, Tbilisi, Georgia; Institute of Archaeology, The Hebrew University of Jerusalem, Jerusalem, Israel; Israel Antiquities Authority, Jerusalem, Israel; Zinman Institute of Archaeology, University of Haifa, Haifa, Israel; CIAS, Department of Life Sciences, University of Coimbra, 3000-456 Coimbra, Portugal; Department of Human Evolution, Max-Planck-Institute for Evolutionary Anthropology, Leipzig, Germany

## Abstract

Archaeological sediments have been shown to preserve ancient DNA, but so far have not yielded genome-scale information of the magnitude of skeletal remains. We retrieved and analysed human and mammalian low-coverage nuclear and high-coverage mitochondrial genomes from Upper Palaeolithic sediments from Satsurblia cave, western Georgia, dated to 25,000 years ago. First, a human female genome with substantial basal Eurasian ancestry, which was an ancestry component of the majority of post-Ice Age people in the Near East, North Africa, and parts of Europe. Second, a wolf genome that is basal to extant Eurasian wolves and dogs and represents a previously unknown, likely extinct, Caucasian lineage that diverged from the ancestors of modern wolves and dogs before these diversified. Third, a bison genome that is basal to present-day populations, suggesting that population structure has been substantially reshaped since the Last Glacial Maximum. Our results provide new insights into the late Pleistocene genetic histories of these three species, and demonstrate that sediment DNA can be used not only for species identification, but also be a source of genome-wide ancestry information and genetic history.

**Highlights:**

- We demonstrate for the first time that genome sequencing from sediments is comparable to that of skeletal remains
- A single Pleistocene sediment sample from the Caucasus yielded three low-coverage mammalian ancient genomes
- We show that sediment ancient DNA can reveal important aspects of the human and faunal past
- Evidence of an uncharacterized human lineage from the Caucasus before the Last Glacial Maximum
- ∼0.01-fold coverage wolf and bison genomes are both basal to present-day diversity, suggesting reshaping of population structure in both species

## Introduction

Ancient genome sequencing from bone (Hagelberg et al., 1989), teeth (Höss et al., 1996) and hair (Gilbert et al., 2007) has revolutionised our understanding of natural history and the human past (Hofreiter et al., 2001; Racimo et al., 2020). When skeletal material is not available, sediment ancient DNA has sometimes been used to determine the presence or absence of different species. Several studies have demonstrated the presence of ancient DNA in sediments, initially with PCR-based methods (Willerslev et al., 2003) and more recently using high throughput sequencing techniques (Pedersen et al., 2016; Søe et al., 2018; Willerslev et al., 2014). Sediment DNA has also been used to track the presence of absence of species across a range of environments and time periods, primarily through targeted amplification or capture of single genetic regions (Pedersen et al., 2015). A ground-breaking study showed DNA preservation in clay-rich sediments for at least 240 ky. (Slon et al., 2017), and used targeted enrichment to recover sufficient numbers of fragments to reconstruct mtDNA phylogenies of Neanderthals and Denisovans. A recent study similarly recovered Denisovan mitochondrial DNA from sediments deposited ∼100 kya and ∼60 kya from Baishiya Karst Cave (BKC) on the Tibetan Plateau (Zhang et al., 2020).

However, the use of sediment DNA has thus far remained limited to species identification or mitochondrial capture. A major question, which we address here, is whether it can also be leveraged to provide genome-wide data informative about overall genetic ancestry and other biological properties of the organisms it derives from such as: sex, presence of multiple individuals or differnetial ancestry clusters. We report results from shotgun sequencing of a sediment sample from the Upper Paleolithic site of Satsurblia Cave, southern Caucasus, that dates to the Last Glacial Maximum (LGM, 25 thousands years ago [kya]). In most of the Caucasus and particularly in western Georgia, karst systems hold low and stable year-round temperatures and low acidity (no guano deposits in most systems), conditions which are particularly favourable to ancient DNA survival. A single sample yielded up to several million sequence reads from human, wolf (*Canis lupus*) and bison (*Bison*), corresponding to genome-wide data comparable to low-coverage sequencing obtained from skeletal remains.s

## Results

We analyzed six sediment samples from different layers of areas A (pre-LGM) and B (pre and post-LGM) of Satsurblia cave (Figure1A) (Pinhasi et al., 2014), and performed sequencing to screen them for mammalian DNA (Table S1). One of the samples, SAT16 LS29 (SAT29), from layer BIII (Figure 1B), which is radiocarbon dated to 25.4-24.5 ka calBP (Pinhasi et al., 2014), contained substantial amounts of DNA from humans as well as from other mammals, and was therefore sequenced to greater depth.

**Figure 1:**
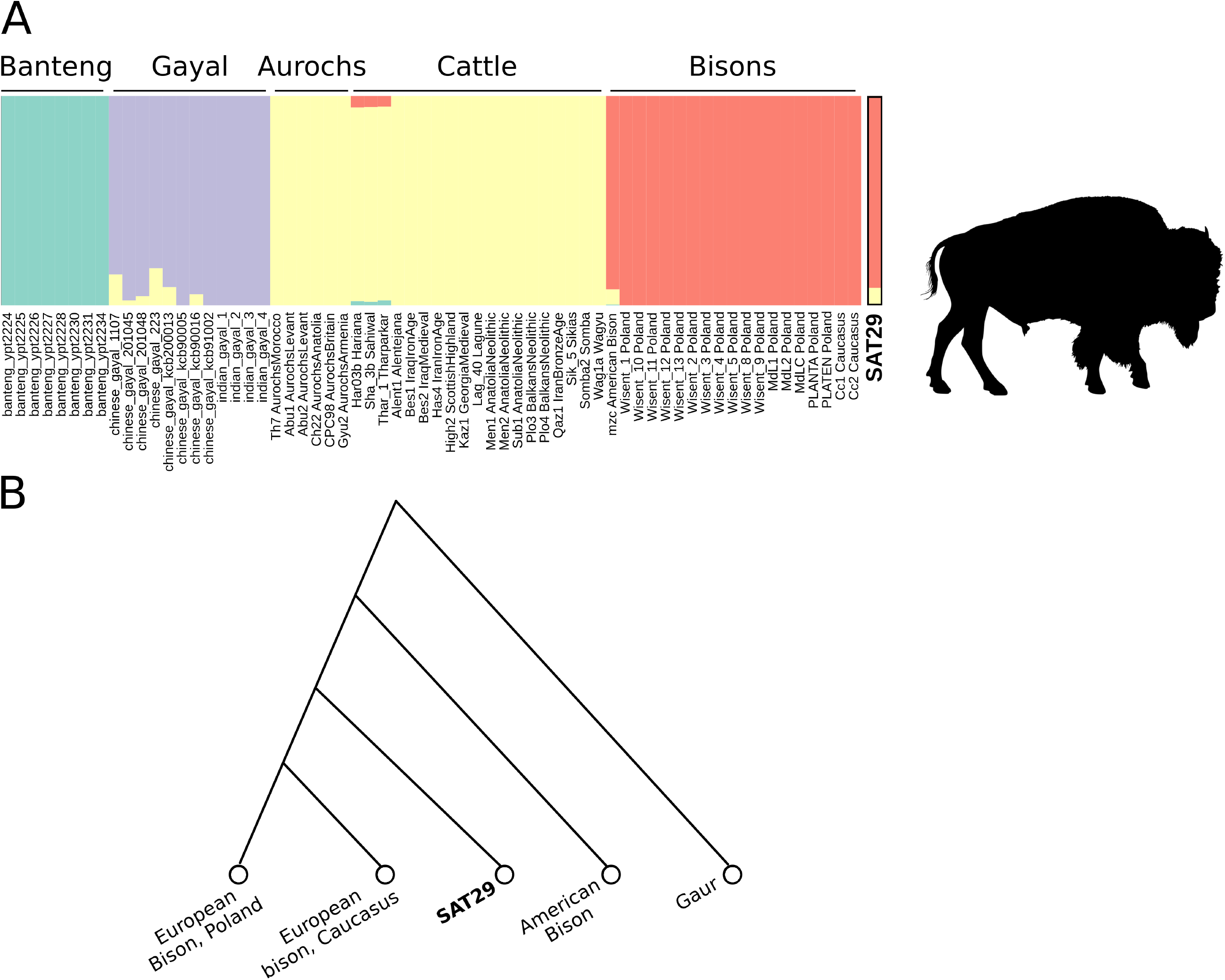
An overview of the location of Satsurblia cave with contextual information. A) Map of the Caucasus region showing relevant sites that have yielded ancient DNA from humans (blue dots), animals (red dot) or both (purple dot). Only sites with remains from the Mesolithic or older are shown. B) Layer B of Satsurblia cave with the SAT29 sampling location shown.

Metagenomic screening of SAT29 performed with Centrifuge (Kim et al., 2016) showed the presence of four mammalian genera among the eukaryotic reads: *Ovis* (28%), *Homo* (9%), *Canis* (5.5%) and *Bos* (2.1%) (Figure S1A). After competitive mapping to the sheep, human, dog and domestic cattle reference genomes (Method Details), we assigned a total of 4,956,676 reads as follows: *Canis* (2,378,237 reads, 48.0% of assigned reads), *Bos* (1,811,555 reads, 36.5%), *Homo* (661,765 reads, 13.5%) and *Ovis* (105,119 reads, 2.1%). Reads mapping to all four species have fragment lengths and deamination patterns typical of ancient DNA (Figure S1B-S1C).

Using the 661,765 human reads, we genotyped 11,116 pseudo-haploid positions from the 1240K dataset (Haak et al., 2015). To explore the human ancestry of SAT29 within the context of pre and post-LGM diversity we performed a Principal Components Analysis (PCA) on 2,335 modern Eurasian genomes (Table S2), and projected 78 ancient individuals onto the resulting components (Figure 2A, Table S2). Previous paleogenomic studies revealed two different ancient human lineages from the Caucasus that were distinct from the rest Pleistocene and early-Holocene diversity. A late Upper Palaeolithic (13.3 kya) genome from Satsurblia cave and a Mesolithic (9.7 kya) genome from the cave of Kotias Klde revealed what has been termed Caucasus Hunter Gatherer (CHG) ancestry, a distinct ancient clade that split from western hunter-gatherers ∼45 kya shortly after the expansion of anatomically modern humans into Western Eurasia (Jones et al., 2015). A second, older, pre-LGM lineage, represented by genome-wide data from two individuals dated to ∼26 kya from Dzudzuana cave, southern Caucasus, is associated with the peoples of the Near East and North Africa, and accounts for at least 46% of their ancestries (Lazaridis et al., 2018). We find that the SAT29 sample clusters with the Dzudzuana2 individual and not with the late Upper Paleolithic and Mesolithic genomes from the region, or with any of the other pre-LGM Eurasian genomes. Unsupervised ADMIXTURE clustering (Alexander and Lange, 2011) further supports a high similarity between SAT29 and Dzudzuana2 (Figure 2B, Figure S2).

**Figure 2:**
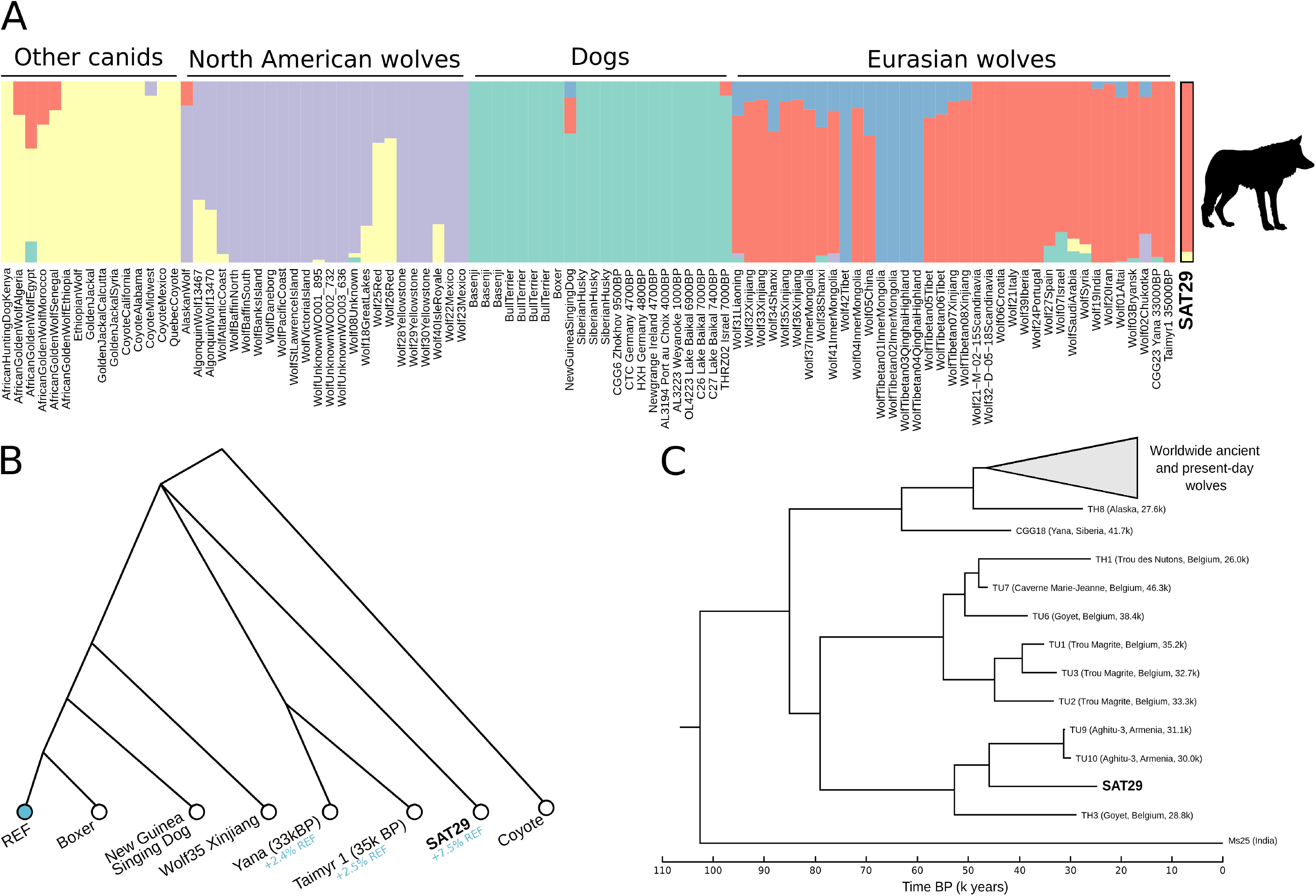
SAT29 human genomics. A) A principal component analysis built with 2,335 modern Eurasian samples into which 78 ancient samples were projected. SAT29 appears closest to Dzudzuana2. B) In ADMIXTURE clustering with 78 ancient genomes, SAT29 displays a profile similar to that of Dzudzuana2. C) Maximum Parsimony tree built with 67 ancient and 168 modern Eurasian mitochondrial genomes. SAT29 falls close to the Upper Paleolithic samples from Bacho Kiro cave in Bulgaria.

We used outgroup *f*_3_-statistics to quantify the amount of shared genetic drift between SAT29 and other ancient genomes (Patterson et al., 2012). SAT29 shares more drift with Villabruna (Italy, 12140±70 bp) (Fu et al., 2016) and Dzudzuana2 than with other ancient individuals (Figure S3A), including the post-LGM individuals from the Caucasus (Satsurblia and Kotias). Among present-day Eurasian populations, SAT29 shows higher genetic affinity to Northern and Western Europeans rather than Central and Southern Asians (Figure S3B).

Through targeted capture (Maricic et al., 2010) we recovered 4,953 human mitochondrial DNA (mtDNA) reads, providing 15-fold coverage of the mtDNA. The consensus sequence has 16 derived positions (Table S3) compared to the rCRS sequence and belongs to haplogroup N like individual Dzudzuana3 from Dzudzuana cave (Figure 2C). We built a maximum parsimony (MP) tree with 68 ancient and 167 modern human mt genomes (Table S4, Figure 2C). The SAT29 sequence is positioned on a branch together with BK-CC7-355 (42450 ± 510 ka) and BK-BB7-240 (41850 ± 480 ka) from the Bacho-Kiro site in Bulgaria, the most ancient west-Eurasian mitochondrial sequences (Hublin et al., 2020) as well as Dzudzuana3 (Lazaridis et al., 2018).

We estimate 1% Neanderthal ancestry in the SAT29 sample, although with large uncertainty due to the low amount of data (95% confidence intervals: 0-6.6%). This point estimate is similar to that of Dzuzuana2 and likely lower than that of Palaeolithic Europeans due to dilution from Basal Eurasian ancestry (Lazaridis et al., 2018)

Our results from the human genomic data from the SAT29 sample are thus consistent with the study by Lazaridis et al (2018), revealing a a previously undocumented pre-LGM human ancestry from the Caucasus that contributed to various succeeding Eurasian populations. The low coverage of the SAT29 genome, however, did not allow us to detect any differences in ancestry patterns between Dzuzuana2 and SAT29.

We analyzed the 2,378,237 *Canis* reads using a set of variants among 722 modern wolves, dogs and other canid species (Plassais et al., 2019). We randomly sampled one SAT29 read at each position, resulting in a genotype call at 439,426 transversion variants. Using ADMIXTURE clustering (Figure 3A) and *f*_*4*_-statistics, we found that the SAT29 sample clearly shares genetic drift with wolves and dogs, to the exclusion of coyotes, golden jackals and other canids (Z > 30 for all species, Method Detail). It does not, however, display stronger affinity to wolves over dogs, or vice versa (Figure S4).

**Figure 3:**
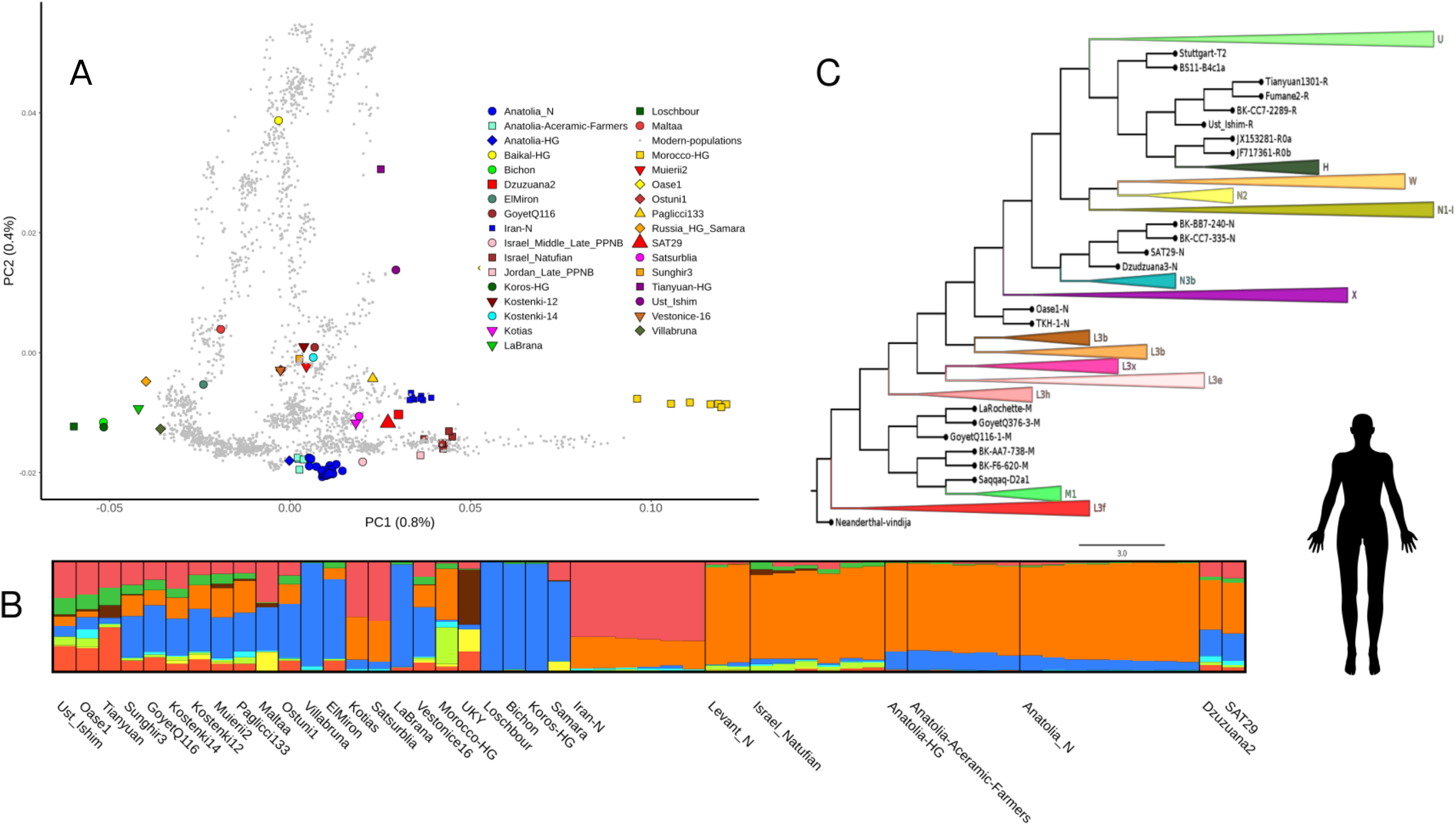
SAT29 wolf genomics. A) In ADMIXTURE clustering with wolves, dogs and other canid species, the SAT29 sample clusters with Eurasian wolves. B) Population history model relating the SAT29 sample to modern wolves, dogs and Pleistocene Siberian wolves. Inferred admixture proportions from the dog reference genome (REF), to account for reference bias, are shown in blue. The trifurcation point indicates that it cannot confidently be determined on which side of this point the SAT29 sample falls. C) Bayesian tree of 68 modern and 39 ancient wolf mitochondrial genomes.

To further investigate the relationship of the SAT29 canid to wolves and dogs, we incorporated two Pleistocene wolf genomes from Siberia (35-33 ka), which have ancestries that are basal to modern wolves and dogs (Sinding et al., 2020; Skoglund et al., 2015). We tested all possible topologies without admixture relating a coyote, SAT29, a modern wolf, a modern dog and the two Pleistocene Siberian wolves, while explicitly accounting for reference bias in the ancient genomes (Method Details). Three of the 100 graphs provide good fits, and feature the Siberian Pleistocene wolves on a branch basal to modern populations. The graphs fit equally well and differ only in that SAT29 is placed either basal to the Siberian Pleistocene branch, on this branch, or downstream of this branch (Figure 3B). Previous studies have found that present-day wolf population structure has mostly formed after the LGM (Freedman et al., 2014; Skoglund et al., 2015; Fan et al., 2016; Loog et al., 2020; Ramos-Madrigal et al., 2020). Our results are consistent with this scenario, as the SAT29 canid harboured an ancestry that diverged from the ancestors of modern wolves and dogs before these diversified. While Late Pleistocene wolves in the Caucasus were not closely related to those in Siberia, they thus had a similarly basal ancestry that has either gone extinct or been overwritten by later population processes.

Through targeted capture (Maricic et al., 2010) we recovered 9,884 *Canis* mtDNA reads, providing 13-fold coverage of the mtDNA. The SAT29 canid consensus sequence falls on a branch together with two ancient wolves from the Aghitu-3 cave in Armenia (Figure 3C), dated to 31-30 ka, on a seemingly extinct West Eurasian branch of the wolf mtDNA phylogeny (Loog et al., 2020).

We inferred the tip date of both the SAT29 human and wolf consensus mtDNA with various priors and datasets (Method Details). The human SAT29 mtDNA consensus has a mean age of 38,815 - 48,243 years bp (95% CI: 19,772 - 74,475 years bp) (Table S5) while the wolf mtDNA consensus mean age is 22,991 - 31,416 years bp (95% CI: 11,712 - 49,884 years bp) (Table S5). The direct date of the BIII layer of Satsurblia cave, from which SAT29 was taken, is 25.5.-24.2 ka cal. bp, and thus falls within the confidence intervals of the mtDNA tip dates. While the mean estimate for the human sequence is older than for the wolf, the 95% confidence intervals overlap across all combinations of datasets and priors used except one.

We compiled a number of Bovinae genomes (Verdugo et al., 2019; Wecek et al., 2016; Wu et al., 2018), identified 1.4 million heterozygous transversion sites in a Gaur genome, and assigned genotypes to all individuals at these sites by randomly sampling one read per position. This resulted in a genotype call at 27,724 transversion variants for the SAT29 bovid sample. Using ADMIXTURE clustering (Figure 4A) and *f*_4_-statistics, we find that the SAT29 bovid shares genetic drift with bisons (*Bison sp*.), to the exclusion of aurochs, domestic cattle and Asian bovid species (Z > 20 for all species, Method Details). This provides important authentication of the soil metagenomic approach, since the identified species is not the cattle (*Bos taurus*) reference genome. To explore the ancestry of the SAT29 bovine reads, we incorporated genomes from early 20th century European bison (*Bison bonasus;* wisents) from Poland and the Caucasus. SAT29 is closer to these, as well as to modern European bison, than to a North American bison (|Z| > 6), implying that the divergence between European and American populations predates the age of SAT29. The Caucasian and the Polish populations have been classified as separate subspecies of the European bison, but the SAT29 sample is not detectably closer to one over the other. Furthermore, these two recent populations share genetic drift to the exclusion of the SAT29 sample (|Z| = 3.4 and 4.5 for the two relevant *f*_4_-statistics).

**Figure 4:**
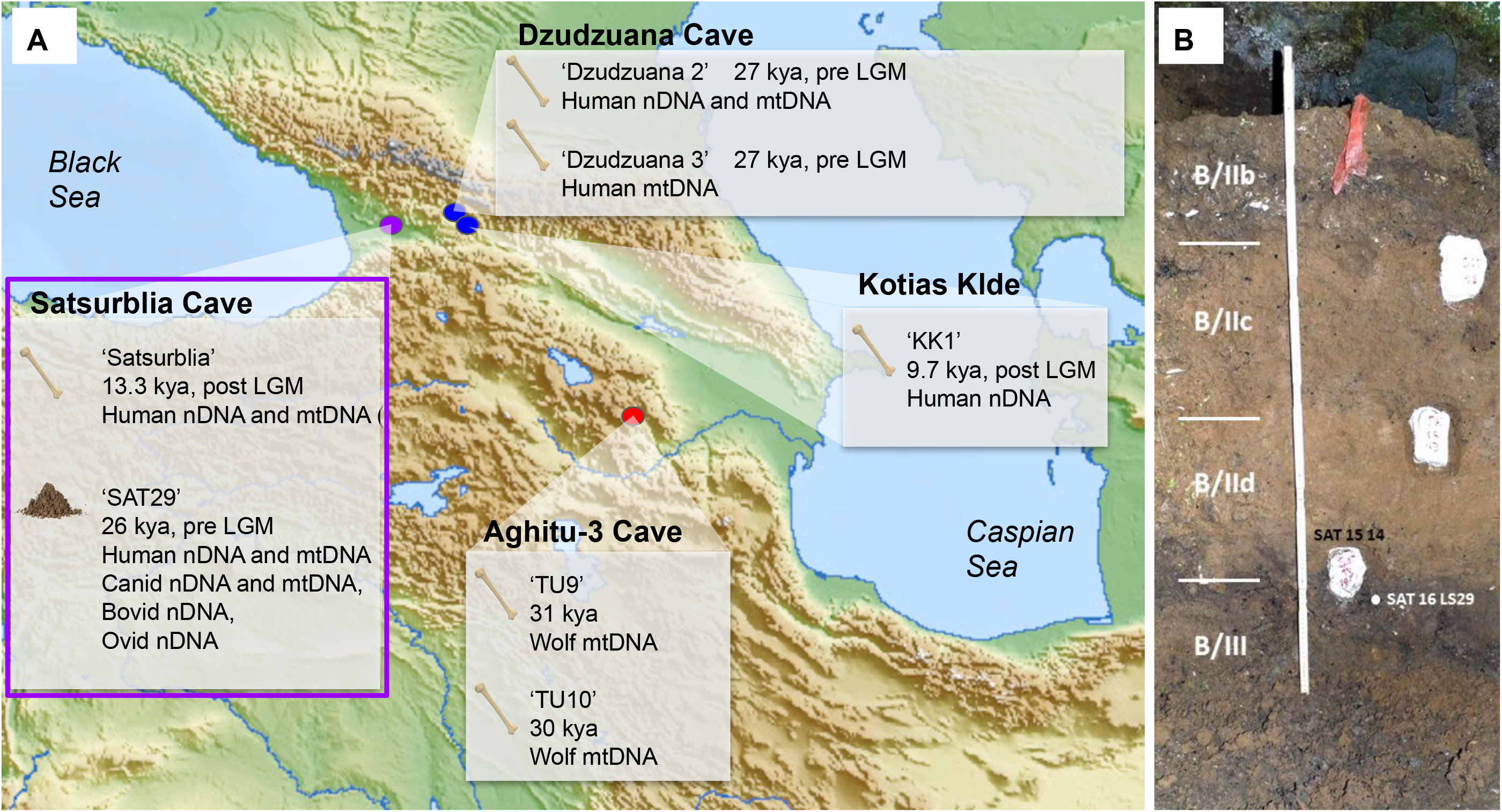
SAT29 bison genomics. A) In ADMIXTURE clustering with cattle, aurochs, bisons, gayals and bantengs, the SAT29 sample clusters with bisons. B) A population history model relating the SAT29 sample to present-day American and historical (early 20th century) European bisons from Poland and the Caucasus.

We tested all possible topologies to summarize the relationships between these bison genomes. The best-fitting topology has SAT29 as basal to the historical genomes from Poland and the Caucasus, and the American bison as basal to all of these (Figure 4). This model explains all *f*_*4*_-statistics except the aforementioned signal of some excess affinity between the American and Polish genomes. We do not attempt to discriminate between more complex models arising if we allow additional admixture events.Overall, the results tentatively suggest that the history of bisons in western Eurasia might share some features with wolf history, in that late Pleistocene ancestries appear basal to present-day populations, suggesting that population structure has been substantially reshaped since the LGM.

We additionally recovered 72,100 reads aligning to *Ovis aries* and explored these reads with a dataset of 103 *Ovis* and *Capra* genomes from 10 species. Tests of the form *f*_*4*_(Oreamnos,SAT29;C,D) indicated that the SAT29 sample is more similar to *Ovis* than to *Capra*, but it does not cluster with any of present-day *Ovis* species (Figure S5). We were thus unable to elaborate on the ancestry of this *Ovis* DNA.

Finally, we explored whether the genomic data retrieved from SAT29 could reveal more about the identity of the individuals from which it derived. First, we examined the human and canid mtDNA to ascertain if multiple individuals contributed DNA. An absence of genetic variation among the human mitochondrial reads is consistent with these deriving from a single individual (Method Details), or from multiple individuals carrying the same mitochondrial haplotype. In contrast, we find evidence of variation among the wolf mitochondrial reads, suggesting they derive from multiple individuals (Method Details). Second, we examined the sex of the human, canid and bovid DNA. When comparing the amount of reads mapping to the X-chromosome relative to the autosomes, we find that the human data is consistent with deriving from a female individual. In contrast, the wolf and bison X-chromosome read fractions are intermediate between those expected for male and female karyotypes, suggesting it may derive from individuals of both sexes, again indicating multiple source individuals. Lastly we compared the number of human mitochondrial and nuclear reads in order to assess from which tissue the human DNA came. The log_e_(mt/nuc) coverage of the human genome has a value of 4.85, which according to Furtwängler et al. (Furtwängler et al., 2018) is a value typically associated with samples from petrous bones.

## Discussion

The recovery of genome-wide data from multiple animal species from a single Pleistocene soil sample opens a new avenue compared to what has previously been learned from sediment DNA (Ardelean et al., 2020; Slon et al., 2017; Willerslev et al., 2003; Zhang et al., 2020). Using this data we were able to make inferences about the genetic ancestry and history of three past mammalian populations that lived in the Southern Caucasus ∼25 ka. The damage characteristics of the recovered DNA, the mitochondrial tip dates and the fact that all three genomes represent ancestries that no longer exist among those species demonstrate that the DNA indeed is of Pleistocene origin, and not a product of modern contamination or movement between different soil layers.

The data presented in this study demonstrates that sediment DNA can be used not only to test the presence or absence of species, but also to obtain genome-wide data informative about the ancestry of the individuals from which the DNA derives. While the amount of DNA retrieved here is still much lower than what is possible from some high quality skeletal elements, it provides information comparable to low-coverage ancient genome sequences. The complementary analyses of multiple mammalian species shows the power of this approach to reconstruct numerous population histories. Our results thus expand the possibilities offered by sediment ancient DNA and demonstrate that it can sometimes serve as an alternative source of genome-wide information when skeletal material is not available, potentially even for organisms that do not leave such remains.

## STAR Methods

### Key resources table

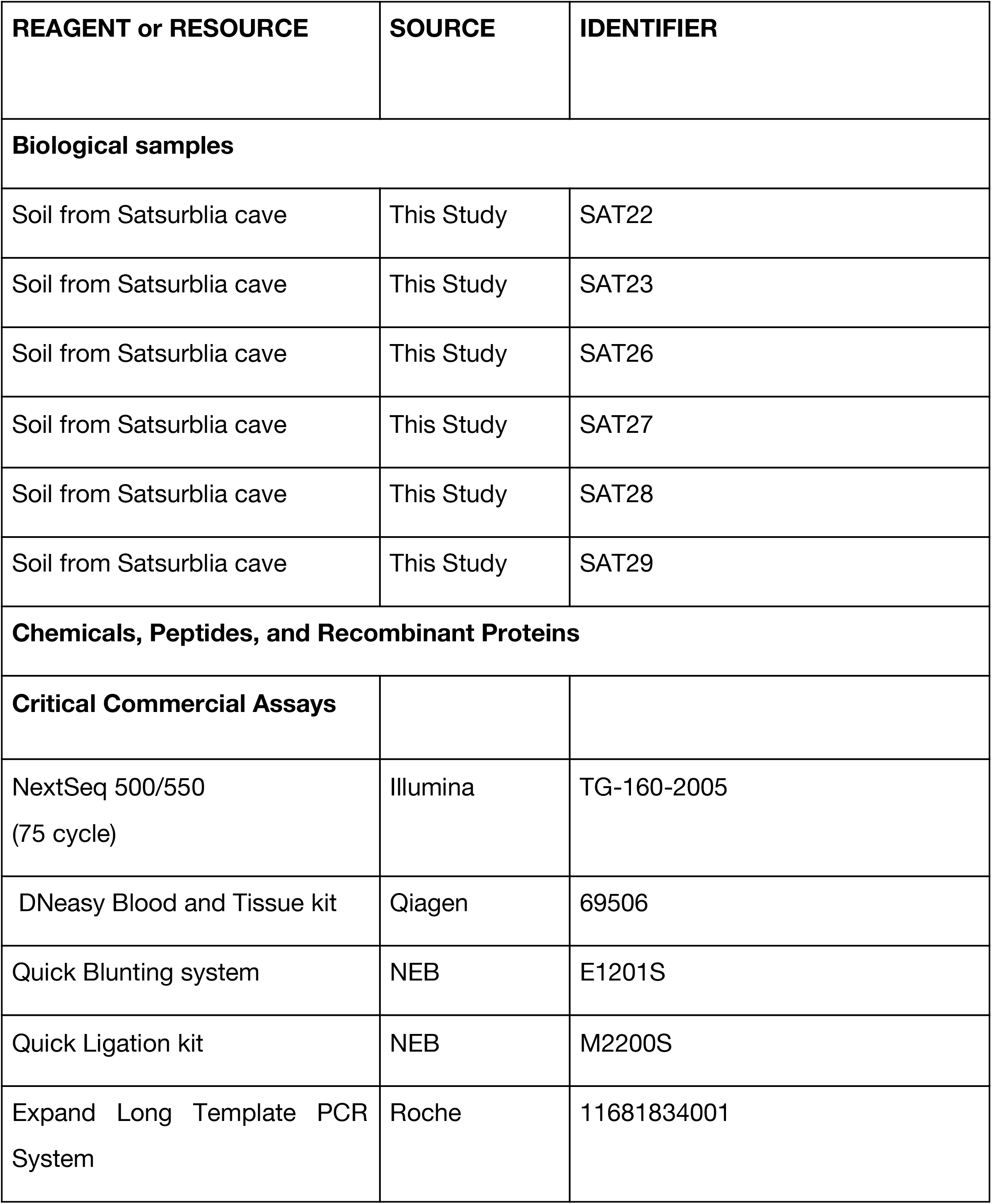
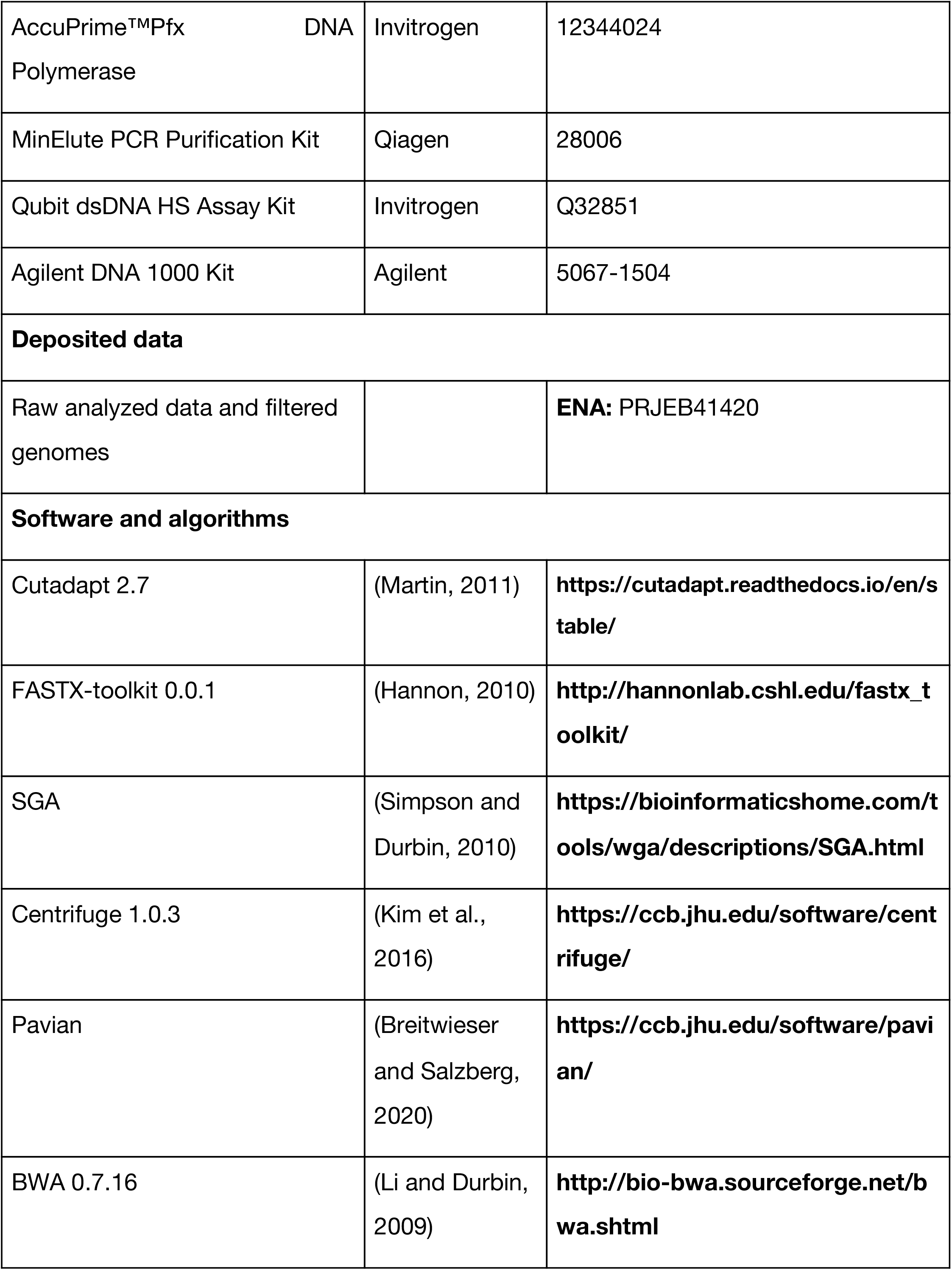
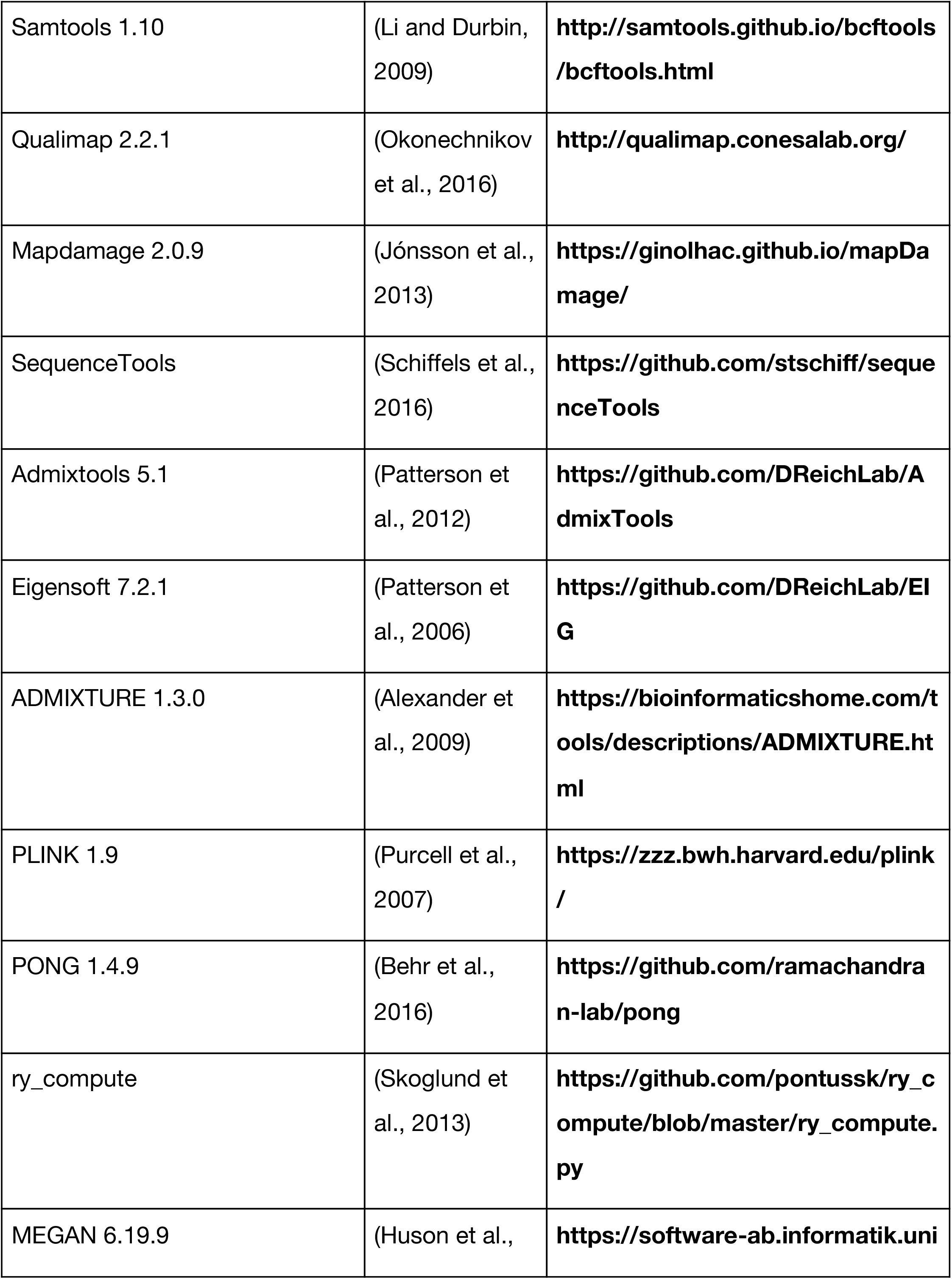
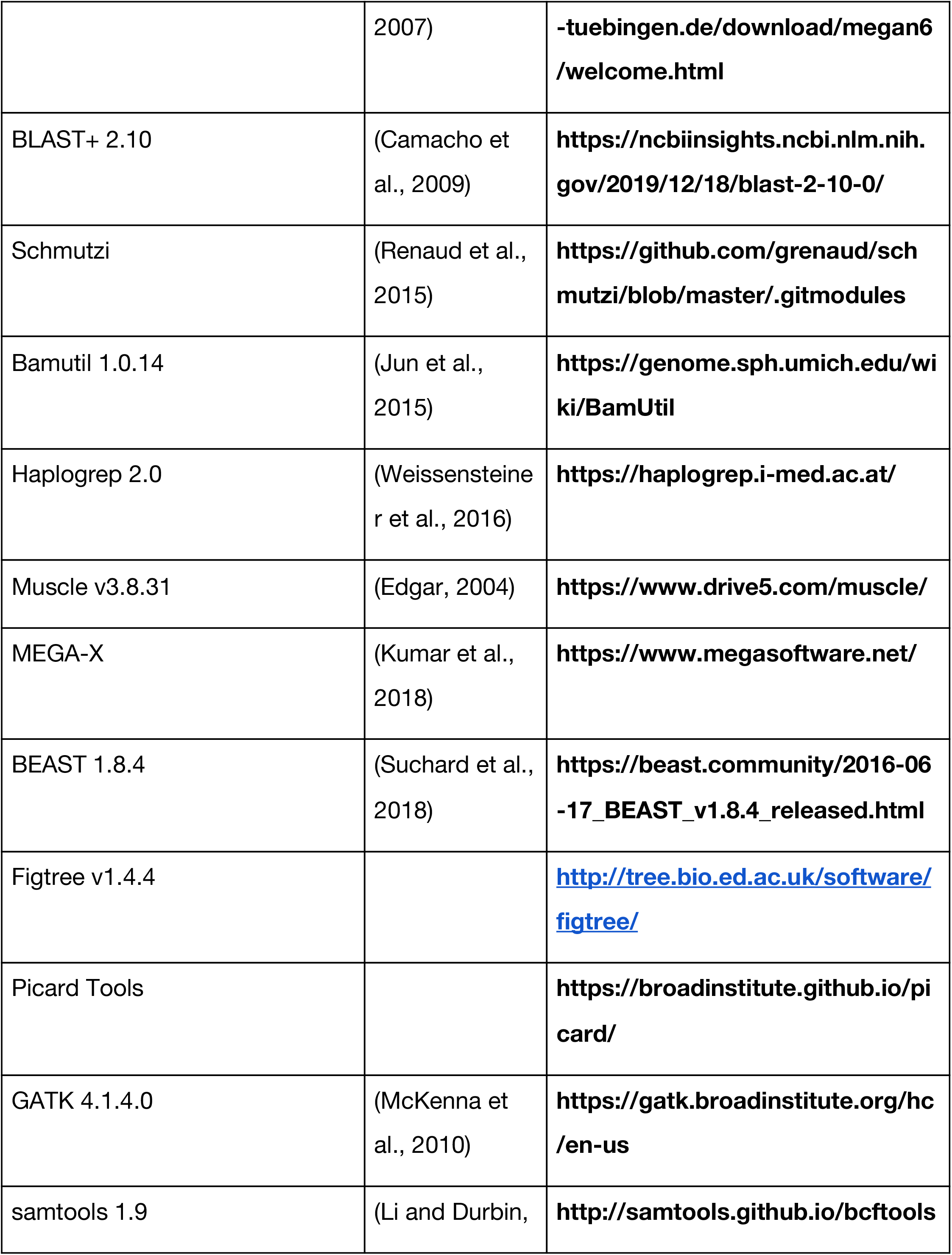
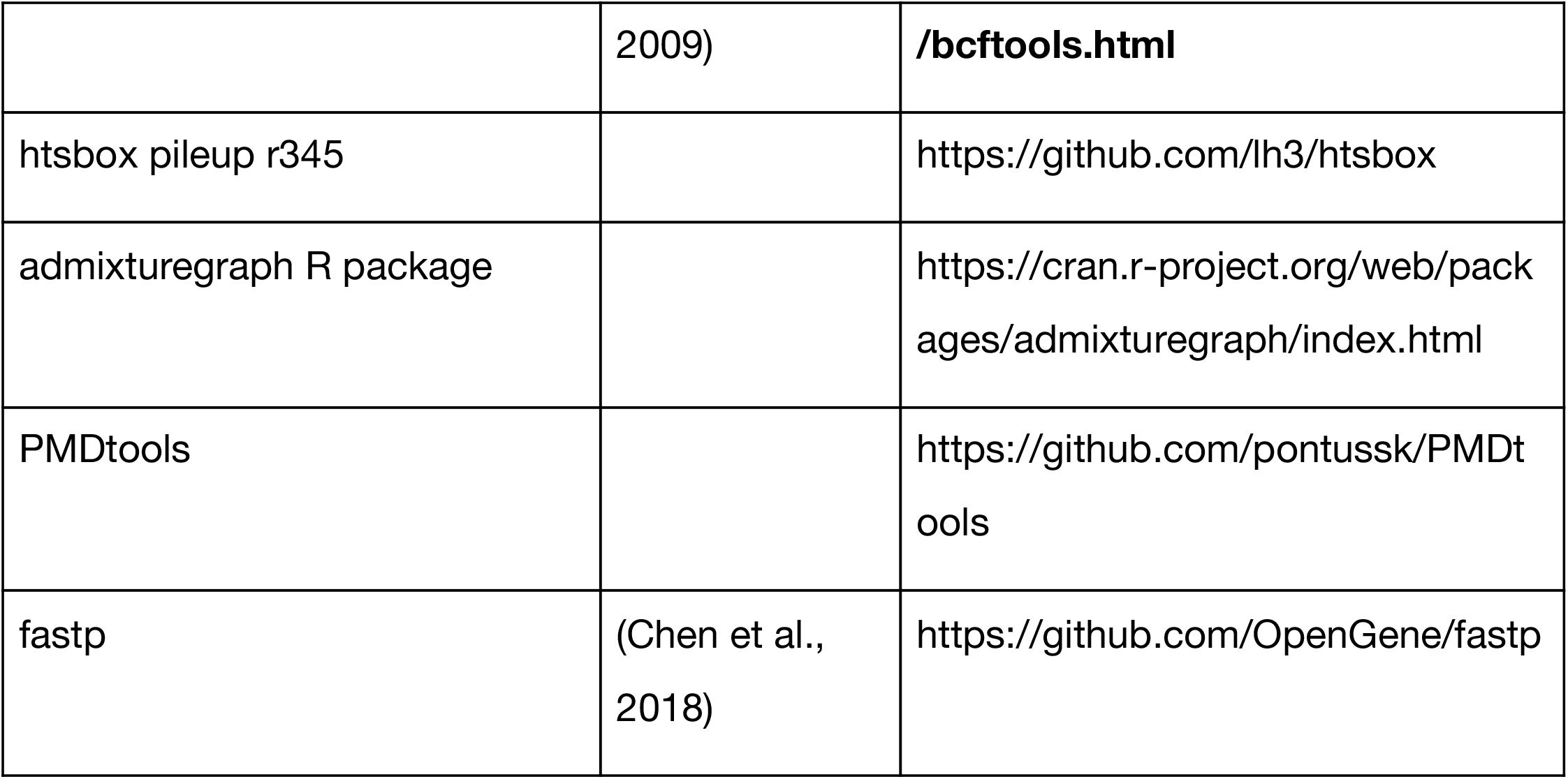

## Resource availability

### Lead contact

Further information on materials, datasets, and protocols should be directed to and will be fulfilled by the Lead Contact, Pere Gelabert

### Data availability

Sequencing data and the filtered genomes are available at the European Nucleotide Archive (ENA) under the accession number: PRJEB41420

## Experimental model and subject details

### Archeological context

Satsurblia Cave is located in Georgia, in the Southern Caucasus region. The cave was discovered in 1975 by N. Kalandadze (Kalandadze and Kalandadze, 1978) and has been excavated during different periods: 1976, 1985–1988, 2008–2010, 2012–2013, and 2016-2017. In the present project we present results from several layers excavated during the 2016 campaign.

Fieldwork in Satsurblia cave has focused on excavations in two areas: Area A in the north-western part of the cave, near the entrance, and Area B in the south-west, in the rear part of the cave. The stratigraphic sequence in Area A contains material remains from the Upper Palaeolithic and Eneolithic periods, and consists, so far, of three main layers: A/I (Eneolithic, 5th millennium BCE), A/II (17.5-16.5 kya), and A/III (25-24 kya). Both areas yielded a series of in situ well preserved Upper Palaeolithic occupational layers and a unique feature of the Upper Palaeolithic layers is the preservation of in situ living surfaces (“floors”), some of them with preserved fireplaces and rich material culture assemblages. The sequence of Area B is divided into five main archaeological layers and encompasses deposits dated to several UP phases: 13 kya (upper part of Layer B/II); 19 kya BP (lower part of Layer B/II), 25-24 kya (lower B/II, B/III, B/IV); and 32-31 kya (Layer B/V) (Pinhasi et al., 2014). A previous genome was published from remains located in square Y5, area A. The radiocarbon dating of this bone is 11,415 ± 50 uncal. BP (OxA-34632) (Jones et al., 2015).

Sediment samples were removed with a knife from the exposed profile sections and in association with block samples and were then stored in zip-up bags and protected from light. Six samples from different layers were sequenced and examined (Table S1). Here we report genomic data retrieved from sediment sample SAT29, which was taken from Area B, Layer BIII, square X4 (Figure 1B). Micromorphological analyses were performed on block sample SAT 15 14, taken next to SAT29, to investigate the formation processes and post-depositional alterations of layer B/III here, and to assess the integrity of the recovered aDNA and their potential source(s). The following presents preliminary results of these analyses (see Figure S6). Natural processes include the weathering of the limestone bedrock with the deposition of limestone clasts and calcareous clay and silt; the deposition of rounded, cross-striated soil aggregates that originate from outside the cave; and the redistribution of clay and their deposition as coatings in large voids by percolating water. Both soil aggregates and clay coatings are connected to repeated water activity leaving the associated clay as an unlikely candidate for lasting DNA absorption in this context. Additionally, soil aggregates show variable color expressions, which result from exposure to heat at different temperatures, again making these aggregates an unlikely source for the recovered aDNA. Heating of soil aggregates and other sediment components results from UP people building combustion features at the site and distributing the combusting residues by trampling and dumping behaviors. Fire use and residues resulting from this behavior - charcoal, ash, burnt sediments and bones - present the dominant anthropogenic component. However, it needs to be noted here that not all bones show heating traces and the heating traces are often limited to charring and low temperature heating. Microscopic bones are sand to gravel size and the most common anthropogenic component in this layer and present a potential source for the recovered aDNA. However, further research into the adsorption and preservation of DNA in archaeological sediments is needed.

## Method Details

### DNA extraction

DNA extraction, library preparation and indexing steps were undertaken in a dedicated aDNA facility within University College Dublin (UCD). All steps were undertaken within a grade B (EU) clean room under grade A unilateral air-flow hoods. Tyvek suits, hair nets, face masks, and nitrile gloves were used to limit contamination. Extraction of soil DNA was performed using 50 mg of soil in an extraction buffer (Collin, 2019) (to final concentration of 0.45M EDTA, 0.02M Tris-HCl (pH 8.0), 0.025% SDS, 0.5mg/mL Proteinase K and dH2O up to final volume). Samples were incubated at 37°C overnight within Matrix E lysing tubes (MPBIO116914) and using an Eppendorf Thermomixer® C with a rotational speed of 1600rpm. Samples were then cleaned according to the method outlined by Dabney et al.(Dabney et al., 2013) and eluted using TET buffer. DNA libraries of the entire extract were prepared using the method outlined by Meyer and Kircher et al. (Meyer and Kircher, 2010) to produce 25uL of the library. Extraction and library negative controls were utilized using 50µl of deionised water.

### PCR, quality control and Next Generation Sequencing

Polymerase Chain Reaction (PCR) amplification and all subsequent steps were undertaken in a grade C laboratory due to increased sample stability. Amplification of 5uL of each library was performed using at a rate of 15 cycles and a single index was added onto the P7 end during amplification (Gamba et al., 2014). The amplified DNA was cleaned using PB and PE buffers (Qiagen 28006). Concentration and molarity (nmol/L) of the working solution were ascertained through Agilent 2100 bioanalyzer and a Qubit4 for fluorometric quantification following manufacturer guidelines. Sequencing was undertaken at UCD Conway Institute of biomolecular and biomedical Research on an Illumina NextSeq 500/550 using the high output v2 (75 cycle) reagent kit (Illumina TG-160-2005). Further sequencing was performed on NovaSeq platforms.

### Mitochondrial Capture

Mitochondrial capture of both the human and canid mtDNA sequences were performed using the method outlined in (Maricic et al., 2010). Briefly, 50uL of modern human or dog blood were used to extract DNA using the Qiagen Blood and Tissue kit. The modern DNA of each species was used for a long-range PCR (Sigma Aldrich Expand™ Long Template PCR System). Two primer pairs were used for the human mtDNA amplification (Gunnarsdóttir et al., 2011) and three primer pairs for the dog mtDNA amplification (Thalmann et al., 2013). The long mtDNA fragments were sheared using a sonicator for eight 15-minute sessions and DNA was checked on a 2% agarose gel to make sure that the DNA was fragmented to below 1Kb in length. Next, fragmented DNA was blunt ended using NEB Quick Blunting system and the BioT/B adapters were ligated to the blunted fragments using the NEB Quick Ligation system to produce the bait for capture.

The single-indexed amplified SAT29 library was re-amplified using Accuprime pfx polymerase and the IS5/IS6 primer pairs (Meyer and Kircher, 2010) for 20 cycles and the concentration was measured on the Qubit 4.0. Before capture, the blocking oligonucleotides BO4, BO6, BO8 and BO10 were used to block the sequencing primer sites. Subsequently the bait and pool were combined and incubated at 65° C for two nights. The enriched DNA was melted off the baits using a 2% NaOH solution and the purified DNA was measured using qPCR to determine the ideal cycle number for amplification. The amplified capture libraries were measured on the Qubit 4.0 and Agilent 2100 bioanalyzer to determine concentration and subsequently sequenced on the Illumina Novaseq system.

### Human DNA screening

We first explored the six libraries for the presence of human DNA. We obtained an average of 14,893,925 reads in each sequencing library. These reads were processed with the methodology described in Collin et al. (Collin et al., 2020). This initial screening showed that five samples exhibited only residual presence of human DNA. The sixth sample, SAT29, had 0.03% of the total reads sequenced map to the human genome. Therefore, this sample was selected for further sequencing. Sequencing and classification results of these samples are presented in Table S1.

### Bioinformatic processing of sample SAT29

After sequencing the SAT29 library to saturation and merging the sequenced reads with the ones from the screening phase, we obtained a total of 561,263,536 reads. These were clipped using Cutadapt 2.7 (Martin, 2011), removing the sequencing adapters and the reads with poly-A tails (reads with more than four As) (Martin, 2011). We also removed reads with qualities below 30 of bases in at least 75% of the read bases with the FASTX-toolkit 0.0.1 (Hannon, 2010). Clipped reads were processed with SGA (Simpson and Durbin, 2010) and redundant reads were removed disabling the kmer check. Finally, two bases per end were trimmed and reads shorter than 30bp were discarded with the FASTX-toolkit (Hannon, 2010). Once collapsed and filtered, we ended with 226,880,778 reads that were used for further analyses. We used Centrifuge 1.0.3 (Kim et al., 2016) with default parameters to classify the sequenced reads into taxa using the whole non-redundant nucleotide database from NCBI indexed following the Centrifuge manual and plotted using Pavian (Breitwieser and Salzberg, 2020). The classification showed the presence of four mammalian taxa with more than 2% of the Eukaryotic classified reads, which we investigated in further analyses: Ovis aries, *Bos* taurus, Homo sapiens and *Canis lupus*. To separate the sequencing reads of the four major mammalian taxa we built a multi-fasta reference file with the genomes of: H. sapiens (GRCh37 Assembly GCA_000001405.1), O. aries (Oar_v3.1, assembly, GCA_000298735.1), B. taurus (ARS-UCD1.2 assembly, GCA_002263795.2) and C. lupus (CanFam3.1 assembly, GCF_000002285.3) following a similar strategy described in Feuerborn et al (Feuerborn et al., 2020). The filtered reads were aligned with bwa aln (Li and Durbin, 2009) disabling seeding, and with a gap open penalty of two. Only reads with mapping qualities above 30 were kept using Samtools 1.10 (Li et al., 2009). Duplicated sequences were removed with picard 2.21.4 ([CSL STYLE ERROR: reference with no printed form.]).. In total 4,956,676 reads were assigned to these species (661,765 *H. sapiens*, 2,378,237 *C. lupus*, 1,811,555 B. taurus and 72,100 O. aries) The characteristics and quality of the mapped reads was assessed with qualimap 2.2.1 (Okonechnikov et al., 2016). We determined the length distribution with fastqc (Andrews, 2010) and assessed the level of damage with mapdamage 2.0.9 (Jónsson et al., 2013). Although two bases per end were clipped the deamination values are notably high: 3’ deamination of 23% in *Bos*, 28% in *Canis*, 20% in *Homo* and 19% in *Ovis* and 5’ deamination of 25% in *Bos*, 30% in *Canis*, 21% in *Homo* and 20% in Ovis. The distribution of these values along the sequence is presented in Figure S3.

### Bacterial analysis

Several bacteria produce collagenase that can degrade collagen and therefore can decompose the bone. These bacteria include the genus: Alcaligenes, Bacillus, Pseudomonas, Clostridium e.g. (Balzer et al., 1997). Additionally it has been shown that the microbial communities change according to the decomposition state of the bones. Skeletal remains tend to have a bacterial profile similar to the soil itself, however several phyla have been seen to be present in archeological remains such as: Pseudomonadaceae, Clostridiaceae, Tissierellaceae, Caulobacteraceae, and Sphingobacteriaceae. (Damann et al., 2015). We have explored such phyla with the results from the Centrifuge classification. The proportion of reads belonging to these phyla showed that: 2,32% of bacterial classified reads correspon to Pseudomonadaceae, 1,02% to *Clostridiaceae, 0*.*45% to Caulobacteraceae and 0*.*11% to Sphingobacteriaceae* In total 4,9% of classified Bacteria are identified as bacteria capable to digest collagen.

### Human population genetics

The final 661,765 filtered human reads were used for the following downstream analyses. We used sequenceTools (Schiffels et al., 2016) to call pseudo-haplotype genotypes of the 1240K dataset (Lazaridis et al., 2016). A total of 11,116 pseudo-haploid positions were recovered. These genotypes were combined with data from 78 ancient genomes (Fu et al., 2014, 2015, 2016; Gamba et al., 2014; Jones et al., 2015; Lazaridis et al., 2016, 2018; van de Loosdrecht et al., 2018; Mathieson et al., 2015; Narasimhan et al., 2019; Olalde et al., 2014; Raghavan et al., 2014; Sikora et al., 2017; Yang et al., 2017a; Yu et al., 2020) (Table S2) and 2,335 present-day individuals from 149 different populations (Jeong et al., 2019; Lazaridis et al., 2014) (Table S2) that were projected on a PCA using eigensoft 7.2.1 (Patterson et al., 2006), using the 597,573 SNPs of the Human Origins (HO) dataset (Lazaridis et al., 2014). We used the option lsqproject in order to minimize the effect of the missing data on the distortion in the PCA location.

Admixture analysis was run using ADMIXTURE 1.3.0 (Alexander et al., 2009) with all individuals from the Human Origins (HO) array and all the available sequences from the David Reich lab database (https://reich.hms.harvard.edu/). The HO dataset SNPs were pruned with option --indep-pairwise of PLINK 1.9 (Purcell et al., 2007) with parameters 250 50 0.4. The total number of remaining SNPs was 436,097. Figure 1C shows the 78 ancient individuals and SAT29 samples with PONG 1.4.9 (Behr et al., 2016).

To explore the genetic affinities and the amount of shared derived SNPs we have used *f*_3_-outgroup statistics using admixtools 5.1 (Patterson et al., 2012) in the form *f*_3_(SAT29,X;Mbuti). X represents both the 78 ancient genomes (Table S2) and the 149 modern populations (Table S2). For the ancient individual comparisons we restricted the analysis to 2,000 shared SNPs and reduced the modern comparisons to 4,000 shared SNPs. We further explored the possible clusterization of SAT29 and Dzuzuana2 individuals with *f*_*4*_ statistics in the form *f*_*4*_(Dzuzuana2,X;SAT29,Mbuti), with X representing the ancient tested populations (Table S6). All these comparisons yielded no concluding results due to the lacking statistical significance due to the low coverage.

In addition, we used qpWave from admixtools 5.1 to test the possible single genetic pool for SAT29 and Dzuduana2. We assigned these two populations as left populations and used Chimp, Altai Neanderthal, Ju_hoan_North, Khomani_San and Vindija as the right populations, from the HO dataset. This yielded to non-significant results (tail probability of. of 0.38).

### Sex determination of SAT29 human reads

For sex determination we used ry_compute (Skoglund et al., 2013). The results show that the SAT29 soil sample is compatible with a female: R_y value of 0.0089 and a CI of: 0.0078-0.0099.

### Neanderthal ancestry in SAT29

We used F4 Ratio (Patterson et al., 2012) to explore the Neanderthal ancestry of the SAT29 sample. We used the genotype data from the 1240k dataset available in (https://reich.hms.harvard.edu/downloadable-genotypes-present-day-and-ancient-dna-data-compiled-published-papers) with the combination: (Chimp AltaiNeandertal: X Mbuti :: Chimp Altai_Neandertal: VindijaNeandertal Mbuti).

### Human mitochondrial analysis

Following the human mtDNA target enrichment step we sequenced 25,483,930 captured reads. After clipping and discarding reads with a base quality score below 30, we had a total of 24,448,710 reads. 2,183,282 reads mapped to the mtDNA human genome. To assure no non-human reads were left after mapping, we used MEGAN 6.19.9 (Huson et al., 2007) and BLAST+ n 2.10 (Altschul et al., 1990) to remove non-human reads aligning all the reads against the whole nt database and selecting only the reads that MEGAN locates in the genus *Homo*. After removing duplicates, our final dataset contained 4,953 reads unique to H. sapiens, which represents 15.31-fold mtDNA genome coverage. The deamination rate of the mtDNA was 0.4 G>A at the 3’ end and 0.41 C>T at the 5’ end.

### mtDNA contamination estimate

The mtDNA coverage for SAT29 is too low to run a standard contamination check with Schmutzi (Renaud et al., 2015). Therefore, we binned our reads using libbam (Renaud, 2018) into three bins using the first and last 10 base pairs to check for deamination: deaminated reads, non-deaminated reads and all reads. We then ran the endocaller script from Schmutzi (Renaud et al., 2015) to compare the reads of the deaminated and non-deaminated bins to look for differences. In the case of contamination levels significant enough to affect consensus calling, we expect to see a difference in the deaminated (ancient reads) and non-deaminated (ancient and potential modern contaminant reads) bins. We only found four positions where the two bins differed and had low consensus conformity. All four positions are a mixture of C and Ts or G and As, indicating that these positions are possibly variable due to deamination, as even the non-deaminated bin of reads could have residual deamination farther into the reads (Table S7). While this method cannot estimate contamination levels directly, the level of contamination is low enough to not influence our consensus calling.

### Presence of multiple individuals

We applied two different methods to ascertain the presence of multiple human individuals in the SAT29 mtDNA. First we used a similar strategy described in Slon et al (Slon et al., 2017). We filtered the reads that showed the presence of deaminated bases in the last five positions on both ends with libbam (Renaud, 2018), and then we filtered for all the positions covered by at least 10 reads. After that we clipped the reads with trimbam from bamutil 1.0.14 (Jun et al., 2015) to minimize the effect of damage. A total of 2,961 positions of the mtDNA positions passed the filters, after which we examined the positions looking for differential base presence. Only 11 positions exhibited the presence of more than two bases different from the majority genotype, but all these positions could be explained by damage that was still present after clipping. Therefore we have not identified positions that could be explained by the presence of multiple individuals. No transversions or substitutions that were not compatible with damage were found.

We then assessed the presence of multiple ancient sequences in the sample using Schmutzi, which indicated the possible presence of another ancient sequence with a percentage of 0.38% (0.355-0.405%). However, a close analysis of the variants revealed that both sequences predicted by Schmutzi, both endogenous and the potential second ancient sequence, only differed in two positions: 310C and 16189C. Both positions are private mutations and supported by less than ⅔ of positions, while the assessment of the mtDNA haplogroup was the same. Therefore we conclude that the results suggest the presence of a single individual or several with the same mitochondrial haplogroup, but these results are inconclusive due to the low coverage.

### Consensus calling

We ran bam2prof and endoCaller scripts from Schmutzi to generate a consensus sequence of the mitochondrial genome. After consensus calling, we selected the positions with a mapping quality higher than 30 and consensus support of more than 67%. The list of derived positions of SAT-29 sequence in respect to the human mtDNA reference genome is presented in Table S3. The total number of positions covered in the consensus is 16,532 out of the total 16,569 positions in the human mtDNA reference genome. The my Haplogroup was assessed with haplogrep

We aligned the SAT29 consensus sequence to 235 mitochondrial genomes of which 168 are present-day and the rest are from Eurasian ancient samples from different publications (Benazzi et al., 2015; Bollongino et al., 2013; Ermini et al., 2008; Fu et al., 2013a, 2013b, 2014; Gilbert et al., 2008; Günther et al., 2018; Hublin et al., 2020; Jones et al., 2015; Krause et al., 2010; Lazaridis et al., 2014; Posth et al., 2016; Sánchez-Quinto et al., 2012; Vai et al., 2019). The samples were aligned with Muscle v3.8.31 (Edgar, 2004) and a maximum parsimony tree with 1000 Bootstrap repetitions was performed with MEGA-X (Kumar et al., 2018).

### Human mitochondrial tip dating

We used six subsets of data for tip dating of the human SAT29 mtDNA: A) A full set of samples (238 samples as shown in Table S4), B) All ancient samples and 14 modern samples that fall close to SAT29 in the maximum parsimony tree as well as the RCRS ancestor (84 samples, Table S4), C) Only ancient samples as well as the RCRS ancestor (70 samples, Table S4), and D) For each of the above sets, a second set was included without the two samples BK-BB7_240 N and BK-CC7-335 N giving a total of six sets.

The full dataset was aligned using Muscle 3.8.31 (Edgar, 2004) and uploaded to MEGA-X where a modeltest was run. The model TN93+G+I had the lowest Bayesian information criterion (Table S8). Each set was then selected and exported as a nexus file. The nexus file was uploaded to BEAUti version 1.10.4. Tip dates were set to the years before the present using the dates shown in Table S4. SAT29 and, if present, the samples BK-BB7_240 N and BK-CC7-335 N were put into their own taxon set and were sampled with individual priors. Each set was run four times with either a strict or an uncorrelated relaxed clock and a coalescent: constant size or coalescent: Bayesian skyline tree prior. SAT29 was given a prior age of 30,000 years BP with a normal distribution, while both BK samples were given a normal distribution prior of 46,000 years BP. BEAST 2.6.0 (Suchard et al., 2018) was run with a 100,000,000 MCMC chain length. The resulting log files were viewed with Tracer v1.7.1 and were checked for ESS above 200. The tree files were annotated with TreeAnnotator v1.10.4 and the resulting annotated trees were viewed with Figtree v1.4.4).

### Determination of the tissue origin

Following the approach described in Furtwängler et al. (Furtwängler et al., 2018) we investigated the most probable tissue of origin of the observed humanDNA in the soil. We divided the mean coverage of the shotgun mtDNA human genome by the mean coverage of the nuclear human genome. We obtained a 1.30-fold mtDNA genome coverage and a 0.019-fold nuclear genome coverage. The loge(mt/nuc) coverage has a value of 4.85, which according to Furtwängler et al. (Furtwängler et al., 2018) is a value typically associated with samples from petrous bones as these bones exhibit mt/nc with values lower than five.

### Wolf genome: comparative dataset

To construct a dataset for ancestry analyses of the SAT29 reads of canid origin, we started from a previously compiled variant call set encompassing 722 dogs, wolves and other canid species (Plassais et al., 2019). We also incorporated a number of additional wolf and other canid genomes from other publications (Gopalakrishnan et al., 2018; Kardos et al., 2018; Liu et al., 2018; Sinding et al., 2018). These additional genomes were mapped to the dog reference genome using bwa mem version 0.7.15 (Li, 2013), marked for duplicates using Picard Tools version 2.18.12 (http://broadinstitute.github.io/picard/), genotyped at the sites present in the 722 canid variant call set using GATK HaplotypeCaller v3.6 (McKenna et al., 2010) with the “-gt_mode GENOTYPE_GIVEN_ALLELES” argument, and then merged into the variant call set using bcftools merge (http://www.htslib.org/).

The variants and genotypes were then filtered by excluding sites displaying excess heterozygosity (“ExcHet” annotation p-value < 1×10−6, as computed using the bcftools fill-tags plugin), setting to missing any genotypes that included an indel allele or any allele with a frequency lower than the two most common alleles at the site and thereby removing such alleles (thus retaining only two SNP alleles overlapping any given position), setting genotypes to missing if the depth at the site (computed as the sum of the “AF” fields) was lower than one third of the genome-wide average coverage of the same, or lower than 5, or higher than twice the average, normalizing allele representation using bcftools norm, and finally excluding sites with missing genotypes for 130 or more individuals. This resulted in 65.5 million SNPs, of which 19.2 million are transversions with a minor allele count in the dataset of at least two.

We assigned genotypes to the SAT29 reads that mapped to the dog genome by sampling one random allele at each of these variants using htsbox pileup r345 (https://github.com/lh3/htsbox) and requiring a minimum read length of 35 (“-l 35”), mapping quality of 30 (“-q 30”) and base quality of 30 (“-Q 30”). We also included data from two Pleistocene Siberian wolves, the 35,000 year old Taimyr-1 from the Taimyr peninsula (Skoglund et al., 2015) and the 33,000 year old CGG23 from the Yana RHS site in eastern Siberia (Sinding et al., 2020), and genotyped these in the same way as the SAT29 data. The SAT29 sample obtained genotype calls at 1,532,986 of the total set of SNPs (2.34%), and 439,426 of the transversions (2.28%).

### Wolf genome: ancestry analyses

We calculated all possible *f*_*4*_-statistics involving SAT29 and the publicly available canid genomes using AdmixTools v5.0 (Patterson et al., 2012), using the qpDstat command with the “*f*_*4*_mode: YES” and “numchrom: 38” arguments.

Using *f*_4_-statistics of the form *f*_4_(AndeanFox,SAT29;X,Wolf35Xinjiang) we find that the SAT29 canid data is closer to a member of *Canis lupus*, Wolf35 from Xinjiang, China, than to representatives of Coyote (Z=31.37), Golden Jackal (Z=37.58), African Golden Wolf (Z=31.01) and Dhole (Z=90.20. The strongly positive values in all tests shows that the data is clearly from a member of the wolf/dog species as opposed to any of these other canid species.

We used the admixturegraph R package (Leppälä et al., 2017) to systematically test admixture graphs by fitting them to the *f4*-statistics. We enumerated all possible graphs involving a coyote (Coyote01, California), a modern Eurasian wolf (Wolf35, Xinjiang), a modern dog (New Guinea singing Dog, pooling individuals NewGuineaSingingDog01, NGSD1, NGSD2 and NGSD3), the two Pleistocene Siberian wolves and the SAT29 sample without admixture events. To each of these admixture graphs, we then grafted on the dog reference genome as a clade with the New Guinea singing dog, and then a boxer individual (Boxer01) as a clade with the reference genome. Because the dog reference genome also derives from a boxer, but a different individual, the contrast between these two can serve to quantify reference bias in other genomes. We therefore introduced, in each graph, an admixture event from the reference genome into each of the ancient genomes – this “reference admixture” can then correct for any systematic shifts in allele frequency caused by reference bias in these genomes. Each graph was then fit five times, retaining the fit that achieved the lowest “best_error” score. Out of the 100 possible graphs, three provided good fits to the data, with the difference between them being only the placement of the SAT29 branch as being on, downstream of, or upstream of the Siberian Pleistocene wolf branch. They correctly predict 209 out of the 210 possible *f*_4_-statistics (|Z| < 3) with the minor outlier statistic: *f*_4_(CoyoteCalifornia,NewGuineaSingingDog; Wolf35Xinjiang,CGG23). After these three, the next best graph has 26 outlier statistics.

### Canid bone testing

We screened five samples from different layers and areas of Satsurblia cave for two main reasons to A) compare the bone DNA and the canid DNA from soil, and B) determine the differential capacity to retrieve DNA from different sources of the same geological layer. In the last excavations no other human bones from the Satsurblia cave have been recovered, however several Canis bones have been identified. We extracted DNA and prepared libraries following the same procedure described for soil. We also captured the dog mtDNA with the same strategy previously described. All five libraries had too few reads for analyses, and therefore were discarded in future analyses (Table S9).

### Canid mitochondrial tip dating

We generated a consensus sequence for the SAT29 canid mitochondrial capture data using htsbox pileup with the “-M” argument for majority consensus sequence, restricting to reads with mapping quality ≥ 30, base quality ≥ 30 and read length ≥ 30 and to positions with at least five reads.

We used two subsets of data for tip dating of the canid SAT29 mtDNA: 1. Full set of samples (107 as shown in Table S10) and 2. Only ancient samples (47 samples, Table S10). The full dataset was aligned using Muscle 3.8.31 and uploaded to MEGA-X where a modeltest was run. The model TN93+G had the lowest Bayesian information criterion. Each set was then selected and exported as a nexus file, and was uploaded to BEAUti version 1.10.4. Tip dates were set to the years before the present using the dates shown in Table S5. SAT29 was put into its own monophyletic taxon set and was sampled with individual priors. Each set was run four times with either a strict or an uncorrelated relaxed clock and a coalescent: constant size or coalescent: Bayesian skyline tree prior (Table S5). SAT29 was given a prior age of 30,000 years BP with a normal distribution. BEAST was run with a 100,000,000 MCMC chain length. The resulting log files were viewed with Tracer v1.7.1 and were checked for ESS above 200. The tree files were annotated with TreeAnnotator v1.10.4 and the resulting annotated trees were viewed with Figtree v1.4.4.

### Presence of multiple individuals

We found evidence for polymorphism in the SAT29 wolf mitochondrial sequences, which could suggest that the retrieved DNA originates from more than one individual. The SAT29 consensus sequence falls on a branch together with two pre-LGM Armenian wolves (TU9 and TU10), but on many sites that define this branch SAT29 also displays observations of the reference allele. We summarized this evidence as follows:

1. We aligned the previously published ancient and modern wolf mitochondrial genomes to the dog mitochondrial reference genome using bwa mem 0.7.17 (Li, 2013) with the “-x intractg” argument, and obtained genotypes for them using htsbox pileup.
2. We merged the SAT29 mitochondrial reads obtained from the targeted capture experiment with those obtained from the shotgun sequencing experiment, to achieve a total coverage of 16.6x (mapping quality ≥ 30, base quality ≥ 30, read length ≥ 35).
3. To reduce the impact of ancient DNA damage and sequencing error on the assessment of polymorphism, we restricted the analysis to a set of polymorphic sites ascertained among the previously published wolf mitochondrial genomes. We first identified sites where the two samples CGG18 (Siberia, 41.7k BP) and TH10 (Alaska, 21k BP) carry the same nucleotide, as a rough approximation of the ancestral sequence of the “major clade” of wolf mitochondria to which these two samples, as well as the majority of ancient and present-day sequences, belong. Any sites containing indels were excluded. We then identified 79 sites on which the Armenian TU10 sample carries a different nucleotide from this major clade. This should constitute a set of variants where SAT29 often should carry the TU10 allele due to its shared phylogenetic history, but might carry the major clade allele if there are other sequences in the sample that carry haplotypes from that clade.
4. We then counted the number of alleles in the SAT29 sample matching the “Armenian” allele and the “major clade” allele at each of these ascertained sites, using htsbox pileup (-q30 -Q30 -l 35). Both alleles are observed at most sites. A few sites display only the major clade allele, but this likely reflects more recent, private mutations in the history of TU10 after its divergence from the SAT29 haplotype.
5. Most of the variants displaying evidence of polymorphism are transitions, which are sensitive to ancient DNA deamination. However, we stratified the variants by mutational direction: derived T->C and A->G mutations in the SAT29 and TU10 sequence cannot be caused by deamination in these samples, while C->T and G->A mutations could be. We observed similar levels of polymorphisms within these two classes of variants, suggesting that ancient DNA damage is not driving the observation of multiple alleles at these sites to any greater extent.
6. We restricted the SAT29 sample to reads displaying evidence of ancient DNA deamination damage, using PMDtools (Skoglund et al., 2014) with the “--threshold 3” argument. While the total read counts are reduced, most sites still display both alleles. This suggests that the additional mitochondrial haplotype(s) in the sample is also of ancient origin, rather than representing modern contamination. The results of such a process are displayed in Figure S7

### Bison genome: comparative dataset

To construct a dataset for ancestry analyses of the SAT29 reads of bovid origin, we downloaded raw sequence reads from the European Nucleotide Archive (ENA) from a number of previously sequenced bovid genomes: present-day gaur (Verdugo et al., 2019; Wu et al., 2018), present-day gayal and banteng (Wu et al., 2018), present-day and ancient domestic taurine and zebu domestic cattle (Verdugo et al., 2019), ancient aurochs (Verdugo et al., 2019), American bison (Wu et al., 2018), present-day European bison from Poland (Wecek et al., 2016; Wu et al., 2018), and the historical (early 20th century) European bison from Poland and the Caucasus (Wecek et al., 2016).

We preprocessed the reads from all the samples using fastp (Chen et al., 2018), filtering through the automatic adapter detection and trimming that applied by default, as well as the “ --low_complexity_filter” and “--length_required 30” arguments. For ancient genomes that had been sequenced paired-end, the “--merge” option was applied and only successfully merged read pairs were retained.

We mapped the filtered reads to the domestic cattle reference genome, using bwa mem v0.7.17 (Li, 2013) in paired-end mode for modern genomes and bwa aln v0.7.17 (Li and Durbin, 2009) in single-end mode, with permissive parameters (“-l 16500 -n 0.01”), for the ancient genomes. We assigned read groups according to the library and run information specified in the ENA metadata for each of the studies, merged reads for each sample and sorted using samtools (Li et al., 2009), and marked duplicate reads using Picard MarkDuplicates v2.18.12 (https://broadinstitute.github.io/picard/).

To define a set of variants to use for ancestry analyses, we identified heterozygous sites in the genome of a single, high-coverage gaur, sample Ga5 (Verdugo et al., 2019). Ascertainment in the Gaur outgroup species, which is estimated to have diverged from bisons more than half a million years ago (Wu et al., 2018), should result in variants that behave in an unbiased fashion in ancestry analyses. We called genotypes in Ga5 using GATK HaplotypeCaller v3.6 (McKenna et al., 2010). We then filtered these genotype calls using bcftools (http://www.htslib.org/) to retain only those variants that were SNPs, were located on the 29 autosomal chromosomes, had a heterozygous genotype, had a genotype quality (GQ field) of >30, a depth (sum of AD fields) of more than 15.04 and less than 49.63 (corresponding to 0.5 and 1.65 times the average autosomal coverage of 30.08, respectively). This resulted in 4,930,425 SNPs, of which 1,447,767 are transversions.

We assigned pseudo-haploid genotypes for all the bovid genomes, including Ga5 itself and the SAT29 reads that mapped to the cattle genome, by sampling one random allele at each of these Ga5 ascertained SNPs, using htsbox pileup r345 (https://github.com/lh3/htsbox) and requiring a minimum read length of 35 (“-l 35”), mapping quality of 30 (“-q 30”) and base quality of 30 (“-Q 30”). The SAT29 sample obtained genotype calls at 94,262 of the total set of SNPs (1.91%), and 27,724 of the transversions (1.91%).

### Bison genome: ancestry analyses

We calculated all possible *f*_*4*_-statistics involving the SAT29 sample and the publicly available bovid genomes using AdmixTools v5.0 (Patterson et al., 2012), using the qpDstat command with the “*f*_*4*_ mode: YES” and “numchrom: 29” arguments. Using *f*_4_-statistics of the form *f*_4_(Ga5.Gaurus,SAT29;X,Wisent11) we find that the SAT29 bovid data is closer a Bison individual, Wisent11 from Poland, than to representatives of aurochs (Gyu2, Armenia, Z=20.59), taurine cattle (ScottishHighland, Z=22.73), Zebu cattle (Tharparkar, Z=24.75), banteng (ypt2230, Z=35.96) and gayal (1107, Z=43.76). The strongly positive values in all tests shows that the data is clearly from a member of the Bison species as opposed to any of these other bovid species.

We used the admixturegraph R package (Leppälä et al., 2017) to systematically test admixture graphs by fitting them to the *f*_*4*_-statistics. We enumerated and fit all possible graphs involving a gaur (Ga5), an American bison (mzc), an historical Polish bison (PLANTA), a historical Caucasian bison (Cc1) and the SAT29 sample with up to one admixture event. Each graph was fit five times, retaining the fit that achieved the lowest “best_error” score.

Among the 15 possible topologies that relate these five genomes without any admixture events, the best-fitting graph has the American bison as basal to the European bisons and SAT29, and then SAT29 as basal to the historical Polish and Caucasian bisons. This graph has just one outlier *f*_*4*_-statistic (|Z| < 3), which fails to account for excess affinity between the American and the Polish bison (*f*_*4*_(Gaur,American bison;Polish Bison,SAT29), Z= −4.03). After this, the next two best-fitting graphs differ from the best-fitting topology in that the position of SAT29 is swapped with that of the historical Polish wisent or the historical Caucasian wisent, respectively. These graphs both feature the same three outlier *f*_*4*_-statistics, the first of which is shared by the best-fitting graph above, and the second and third of which fail to account for shared drift between the historical European bisons to the exclusion of SAT29 (*f*_*4*_(Gaur,Causasian bison;Polish bison,SAT29), Z=−4.51, *f*_*4*_(Gaur,Polish bison;Caucasian bison, SAT29), Z=-3.43.*f*_*4*_

Following these, all other graphs without admixture events have seven or more outlier *f*_*4*_-statistics. When allowing for one admixture event, 10 out of the 315 possible graphs fit the data without any outlier statistics. Multiple solutions with quite variable topologies are thus possible, and with the limited data available we do not attempt to discriminate between these.

### Ovis genomic analysis

We explored the possible phylogenetic position of the reads that aligned to the Oar_v3.1 genome within the Ovis and Capra genus. In order to determine the SNPs to compare to SAT29, we built a dataset with individuals from all the available species of the genus *Ovis* and *Capra*: *Ovis vignei* (Nextgen project: https://projects.ensembl.org/nextgen/), *Ovis aries* (Nextgen project: https://projects.ensembl.org/nextgen/), *Ovis canadensis* (Kardos et al., 2015), *Ovis ammon* (Yang et al., 2017b), *Ovis orientalis* (Nextgen project: https://projects.ensembl.org/nextgen/), *Ovis nivicola* (Upadhyay et al., 2020) Capra hircus (Nextgen Project: https://projects.ensembl.org/nextgen/), *C. caucasica (Zheng et al., 2020), C. ibex (Grossen et al., 2020), C. aegagrus* (Alberto et al., 2018) and Oreamnos americanus as an outgroup (Martchenko et al., 2020). We downloaded the available VCF files of the following genomes: 75 *Ovis aries* genomes, 4 *Ovis vignei* genomes, 14 *Ovis orientalis* genomes and one Capra hircus. This dataset consists of 48,870,177 SNPs in the autosomal chromosomes of the *Ovis aries* genome. After filtering SNPs for MAF < 0.05 and removing non-SNPs and no-biallelic SNPs and SNPs not located in autosomes, 22,553,044 SNPs were kept.

As the available VCF files does not cover all species we wanted to include, we downloaded the FASTQ files of one Ovis canadensis, one Ovis nivicola, one Ovis ammon, one C caucasica, one Capra ibex, one C sibirica, two C aegagrus and one Oreamnos americanus to produce additional VCF files for downstream analyses. The sequencing reads of the FASTQ files were aligned with BWA (Li and Durbin, 2009), duplicate reads were removed with picard ([CSL STYLE ERROR: reference with no printed form.]) and low quality reads (<30) were removed with Samtools (Li et al., 2009). These filtered reads were used for variant calling with GATK HaplotypeCaller v3.6 (67) by genotyping the positions of the filtered dataset with the “-gt_mode GENOTYPE_GIVEN_ALLELES” argument. The reads were then merged into the variant call set using bcftools merge (http://www.htslib.org/). Genotypes were filtered with bcftools for: MBQ>30, depth of coverage below the half of the average coverage and more than double of the average coverage to eliminate possible misalignments in low and high complexity regions. These new VCF files were then merged with the downloaded VCF files. The final dataset was filtered for positions with more than 10% of missing sites and excess of Heterozygosity (pval < 1.10-6).

We called pseudo haplotype genotypes using the 22 million positions of SAT29 Ovis reads using Sequence Tools (Schiffels et al., 2016) and recovered 19,469 SNPs of the SAT29 genome. We used *f*_*4*_ statistics from admixtools (Patterson et al., 2012) to determine the closest taxa to the SAT29 sample. The analysis did not yield any concluding result as the number of SNPS is likely too low.

## Acknowledgments

This study makes use of data generated by the NextGen Consortium. We acknowledge Gabriel Renaud for advice on Schmutzi and Spencer Sawyer, Manuela Alscher and Odin for modern DNA.

## Funding

This work has been supported by the “Mineralogical Preservation of the Human Biome from the Depth of Time” (MINERVA) research platform, code AGB326800. P.S. was supported by the European Research Council (grant no. 852558), a Wellcome Trust Investigator award (217223/Z/19/Z), the Vallee Foundation, and Francis Crick Institute core funding (FC001595) from Cancer Research UK, the UK Medical Research Council, and the Wellcome Trust. T.C.C was supported by the Medical Trainee PhD Scholarship, UCD.

## Author contributions

R.P, S.S and P.G conceived the study. A. BC, D.L, T. M, N.J, Z.M and G.B provided samples, M.S collected the samples, S.S, O.C, D.F, T. C. C, V.O K.TO and R. N. M. F. performed the experimental work, P.G, S.S, A.B and T. C. C. analyzed the data. P.G, S.S, A.B, R.P and P.S wrote the manuscript with inputs from all coauthors.

## Notes

### Competing Interest Statement

The authors have declared no competing interest.

## References

Alberto, F.J., Boyer, F., Orozco-terWengel, P., Streeter, I., Servin, B., de Villemereuil, P., Benjelloun, B., Librado, P., Biscarini, F., Colli, L., et al. (2018). Convergent genomic signatures of domestication in sheep and goats. Nat. Commun. 9, 813.

Alexander, D.H., and Lange, K. (2011). Enhancements to the ADMIXTURE algorithm for individual ancestry estimation. BMC Bioinformatics 12, 246.

Alexander, D.H., Novembre, J., and Lange, K. (2009). Fast model-based estimation of ancestry in unrelated individuals. Genome Res. 19, 1655–1664.

Altschul, S.F., Gish, W., Miller, W., Myers, E.W., and Lipman, D.J. (1990). Basic local alignment search tool. J. Mol. Biol. 215, 403–410.

Andrews, S. (2010). A Quality Control Tool for High Throughput Sequence Data.

Ardelean, C.F., Becerra-Valdivia, L., Pedersen, M.W., Schwenninger, J.-L., Oviatt, C.G., Macías-Quintero, J.I., Arroyo-Cabrales, J., Sikora, M., Ocampo-Díaz, Y.Z.E., Rubio-Cisneros, I.I., et al. (2020). Evidence of human occupation in Mexico around the Last Glacial Maximum. Nature 584, 87–92.

Balzer, A., Gleixner, G., Grupe, G., Schmidt, H.-L., Schramm, S., and Turban-Just, S. (1997). IN VITRO DECOMPOSITION OF BONE COLLAGEN BY SOIL BACTERIA: THE IMPLICATIONS FOR STABLE ISOTOPE ANALYSIS IN ARCHAEOMETRY. Archaeometry 39, 415–429.

Behr, A.A., Liu, K.Z., Liu-Fang, G., Nakka, P., and Ramachandran, S. (2016). pong: fast analysis and visualization of latent clusters in population genetic data. Bioinformatics 32, 2817–2823.

Benazzi, S., Slon, V., Talamo, S., Negrino, F., Peresani, M., Bailey, S.E., Sawyer, S., Panetta, D., Vicino, G., Starnini, E., et al. (2015). Archaeology. The makers of the Protoaurignacian and implications for Neandertal extinction. Science 348, 793–796.

Bollongino, R., Nehlich, O., Richards, M.P., Orschiedt, J., Thomas, M.G., Sell, C., Fajkosová, Z., Powell, A., and Burger, J. (2013). 2000 years of parallel societies in Stone Age Central Europe. Science 342, 479–481.

Breitwieser, F.P., and Salzberg, S.L. (2020). Pavian: interactive analysis of metagenomics data for microbiome studies and pathogen identification. Bioinformatics 36, 1303–1304.

Camacho, C., Coulouris, G., Avagyan, V., Ma, N., Papadopoulos, J., Bealer, K., and Madden, T.L. (2009). BLAST+: architecture and applications. BMC Bioinformatics 10, 421.

Chen, S., Zhou, Y., Chen, Y., and Gu, J. (2018). fastp: an ultra-fast all-in-one FASTQ preprocessor. Bioinformatics 34, i884–i890.

Collin, T.C. (2019). Development and Application of a Metagenomic Ancient DNA Approach for the Identification and Assessment of Taxa from Anthropogenic Sediments: Reconstructing the Past. PhD. University College Dublin. Ireland.

Collin, T.C., Drosou, K., O’Riordan, J.D., Meshveliani, T., Pinhasi, R., and Feeney, R.N.M. (2020). An open-sourced bioinformatic pipeline for the processing of Next-Generation Sequencing derived nucleotide reads: Identification and authentication of ancient metagenomic DNA.

Dabney, J., Knapp, M., Glocke, I., Gansauge, M.-T., Weihmann, A., Nickel, B., Valdiosera, C., García, N., Pääbo, S., Arsuaga, J.-L., et al. (2013). Complete mitochondrial genome sequence of a Middle Pleistocene cave bear reconstructed from ultrashort DNA fragments. Proc. Natl. Acad. Sci. U. S. A. 110, 15758–15763.

Damann, F.E., Williams, D.E., and Layton, A.C. (2015). Potential Use of Bacterial Community Succession in Decaying Human Bone for Estimating Postmortem Interval. J. Forensic Sci. 60, 844–850.

Edgar, R.C. (2004). MUSCLE: multiple sequence alignment with high accuracy and high throughput. Nucleic Acids Res. 32, 1792–1797.

Ermini, L., Olivieri, C., Rizzi, E., Corti, G., Bonnal, R., Soares, P., Luciani, S., Marota, I., De Bellis, G., Richards, M.B., et al. (2008). Complete mitochondrial genome sequence of the Tyrolean Iceman. Curr. Biol. 18, 1687–1693.

Feuerborn, T.R., Palkopoulou, E., van der Valk, T., von Seth, J., Munters, A., Pečnerová, P., Dehasque, M., Ureña, I., Ersmark, E., Lagerholm, V.K., et al. (2020). Competitive mapping allows to identify and exclude human DNA contamination in ancient faunal genomic datasets.

Fu, Q., Meyer, M., Gao, X., Stenzel, U., Burbano, H.A., Kelso, J., and Pääbo, S. (2013a). DNA analysis of an early modern human from Tianyuan Cave, China. Proc. Natl. Acad. Sci. U. S. A. 110, 2223–2227.

Fu, Q., Mittnik, A., Johnson, P.L.F., Bos, K., Lari, M., Bollongino, R., Sun, C., Giemsch, L., Schmitz, R., Burger, J., et al. (2013b). A revised timescale for human evolution based on ancient mitochondrial genomes. Curr. Biol. 23, 553–559.

Fu, Q., Li, H., Moorjani, P., Jay, F., Slepchenko, S.M., Bondarev, A.A., Johnson, P.L.F., Aximu-Petri, A., Prüfer, K., de Filippo, C., et al. (2014). Genome sequence of a 45,000-year-old modern human from western Siberia. Nature 514, 445–449.

Fu, Q., Hajdinjak, M., Moldovan, O.T., Constantin, S., Mallick, S., Skoglund, P., Patterson, N., Rohland, N., Lazaridis, I., Nickel, B., et al. (2015). An early modern human from Romania with a recent Neanderthal ancestor. Nature 524, 216–219.

Fu, Q., Posth, C., Hajdinjak, M., Petr, M., Mallick, S., Fernandes, D., Furtwängler, A., Haak, W., Meyer, M., Mittnik, A., et al. (2016). The genetic history of Ice Age Europe. Nature 534, 200–205.

Furtwängler, A., Reiter, E., Neumann, G.U., Siebke, I., Steuri, N., Hafner, A., Lösch, S., Anthes, N., Schuenemann, V.J., and Krause, J. (2018). Ratio of mitochondrial to nuclear DNA affects contamination estimates in ancient DNA analysis. Sci. Rep. 8, 14075.

Gamba, C., Jones, E.R., Teasdale, M.D., McLaughlin, R.L., Gonzalez-Fortes, G., Mattiangeli, V., Domboróczki, L., Kővári, I., Pap, I., Anders, A., et al. (2014). Genome flux and stasis in a five millennium transect of European prehistory. Nat. Commun. 5, 5257.

Gilbert, M.T.P., Tomsho, L.P., Rendulic, S., Packard, M., Drautz, D.I., Sher, A., Tikhonov, A., Dalén, L., Kuznetsova, T., Kosintsev, P., et al. (2007). Whole-genome shotgun sequencing of mitochondria from ancient hair shafts. Science 317, 1927–1930.

Gilbert, M.T.P., Kivisild, T., Grønnow, B., Andersen, P.K., Metspalu, E., Reidla, M., Tamm, E., Axelsson, E., Götherström, A., Campos, P.F., et al. (2008). Paleo-Eskimo mtDNA genome reveals matrilineal discontinuity in Greenland. Science 320, 1787–1789.

Gopalakrishnan, S., Sinding, M.-H.S., Ramos-Madrigal, J., Niemann, J., Samaniego Castruita, J.A., Vieira, F.G., Carøe, C., Montero, M. de M., Kuderna, L., Serres, A., et al. (2018). Interspecific Gene Flow Shaped the Evolution of the Genus Canis. Curr. Biol. 28, 3441–3449.e5.

Grossen, C., Guillaume, F., Keller, L.F., and Croll, D. (2020). Purging of highly deleterious mutations through severe bottlenecks in Alpine ibex. Nat. Commun. 11, 1001.

Gunnarsdóttir, E.D., Li, M., Bauchet, M., Finstermeier, K., and Stoneking, M. (2011). High-throughput sequencing of complete human mtDNA genomes from the Philippines. Genome Res. 21, 1–11.

Günther, T., Malmström, H., Svensson, E.M., Omrak, A., Sánchez-Quinto, F., Kilinç, G.M., Krzewinska, M., Eriksson, G., Fraser, M., Edlund, H., et al. (2018). Population genomics of Mesolithic Scandinavia: Investigating early postglacial migration routes and high-latitude adaptation. PLoS Biol. 16, e2003703.

Haak, W., Lazaridis, I., Patterson, N., Rohland, N., Mallick, S., Llamas, B., Brandt, G., Nordenfelt, S., Harney, E., Stewardson, K., et al. (2015). Massive migration from the steppe was a source for Indo-European languages in Europe. Nature 522, 207–211.

Hagelberg, E., Sykes, B., and Hedges, R. (1989). Ancient bone DNA amplified. Nature 342, 485.

Hannon, G.J. (2010). FASTX-Toolkit.

Hofreiter, M., Serre, D., Poinar, H.N., Kuch, M., and Pääbo, S. (2001). Ancient DNA. Nat. Rev. Genet. 2, 353–359.

Höss, M., Dilling, A., Currant, A., and Pääbo, S. (1996). Molecular phylogeny of the extinct ground sloth Mylodon darwinii. Proc. Natl. Acad. Sci. U. S. A. 93, 181–185.

Hublin, J.-J., Sirakov, N., Aldeias, V., Bailey, S., Bard, E., Delvigne, V., Endarova, E., Fagault, Y., Fewlass, H., Hajdinjak, M., et al. (2020). Initial Upper Palaeolithic Homo sapiens from Bacho Kiro Cave, Bulgaria. Nature 1–4.

Huson, D.H., Auch, A.F., Qi, J., and Schuster, S.C. (2007). MEGAN analysis of metagenomic data. Genome Res. 17, 377–386.

Jeong, C., Balanovsky, O., Lukianova, E., Kahbatkyzy, N., Flegontov, P., Zaporozhchenko, V., Immel, A., Wang, C.-C., Ixan, O., Khussainova, E., et al. (2019). The genetic history of admixture across inner Eurasia. Nat Ecol Evol 3, 966–976.

Jones, E.R., Gonzalez-Fortes, G., Connell, S., Siska, V., Eriksson, A., Martiniano, R., McLaughlin, R.L., Gallego Llorente, M., Cassidy, L.M., Gamba, C., et al. (2015). Upper Palaeolithic genomes reveal deep roots of modern Eurasians. Nat. Commun. 6, 8912.

Jónsson, H., Ginolhac, A., Schubert, M., Johnson, P.L.F., and Orlando, L. (2013). mapDamage2.0: fast approximate Bayesian estimates of ancient DNA damage parameters. Bioinformatics 29, 1682–1684.

Jun, G., Wing, M.K., Abecasis, G.R., and Kang, H.M. (2015). An efficient and scalable analysis framework for variant extraction and refinement from population-scale DNA sequence data. Genome Res. 25, 918–925.

Kalandadze, A.N., and Kalandadze, K.S. (1978). Archaeological Research of Karstic Caves in Tskaltubo region. Caves of Georgia 116–136.

Kardos, M., Luikart, G., Bunch, R., Dewey, S., Edwards, W., McWilliam, S., Stephenson, J., Allendorf, F.W., Hogg, J.T., and Kijas, J. (2015). Whole-genome resequencing uncovers molecular signatures of natural and sexual selection in wild bighorn sheep. Mol. Ecol. 24, 5616–5632.

Kardos, M., Åkesson, M., Fountain, T., Flagstad, Ø., Liberg, O., Olason, P., Sand, H., Wabakken, P., Wikenros, C., and Ellegren, H. (2018). Genomic consequences of intensive inbreeding in an isolated wolf population. Nat Ecol Evol 2, 124–131.

Kim, D., Song, L., Breitwieser, F.P., and Salzberg, S.L. (2016). Centrifuge: rapid and sensitive classification of metagenomic sequences. Genome Res. 26, 1721–1729.

Krause, J., Fu, Q., Good, J.M., Viola, B., Shunkov, M.V., Derevianko, A.P., and Pääbo, S. (2010). The complete mitochondrial DNA genome of an unknown hominin from southern Siberia. Nature 464, 894–897.

Kumar, S., Stecher, G., Li, M., Knyaz, C., and Tamura, K. (2018). MEGA X: Molecular Evolutionary Genetics Analysis across Computing Platforms. Mol. Biol. Evol. 35, 1547–1549.

Lazaridis, I., Patterson, N., Mittnik, A., Renaud, G., Mallick, S., Kirsanow, K., Sudmant, P.H., Schraiber, J.G., Castellano, S., Lipson, M., et al. (2014). Ancient human genomes suggest three ancestral populations for present-day Europeans. Nature 513, 409–413.

Lazaridis, I., Nadel, D., Rollefson, G., Merrett, D.C., Rohland, N., Mallick, S., Fernandes, D., Novak, M., Gamarra, B., Sirak, K., et al. (2016). Genomic insights into the origin of farming in the ancient Near East. Nature 536, 419–424.

Lazaridis, I., Belfer-Cohen, A., Mallick, S., Patterson, N., Cheronet, O., Rohland, N., Bar-Oz, G., Bar-Yosef, O., Jakeli, N., Kvavadze, E., et al. (2018). Paleolithic DNA from the Caucasus reveals core of West Eurasian ancestry. bioRxiv 423079.

Leppälä, K., Nielsen, S.V., and Mailund, T. (2017). admixturegraph: an R package for admixture graph manipulation and fitting. Bioinformatics 33, 1738–1740.

Li, H. (2013). Aligning sequence reads, clone sequences and assembly contigs with BWA-MEM.

Li, H., and Durbin, R. (2009). Fast and accurate short read alignment with Burrows-Wheeler transform. Bioinformatics 25, 1754–1760.

Li, H., Handsaker, B., Wysoker, A., Fennell, T., Ruan, J., Homer, N., Marth, G., Abecasis, G., Durbin, R., and 1000 Genome Project Data Processing Subgroup (2009). The Sequence Alignment/Map format and SAMtools. Bioinformatics 25, 2078–2079.

Liu, Y.-H., Wang, L., Xu, T., Guo, X., Li, Y., Yin, T.-T., Yang, H.-C., Hu, Y., Adeola, A.C., Sanke, O.J., et al. (2018). Whole-Genome Sequencing of African Dogs Provides Insights into Adaptations against Tropical Parasites. Mol. Biol. Evol. 35, 287–298.

Loog, L., Thalmann, O., Sinding, M.-H.S., Schuenemann, V.J., Perri, A., Germonpré, M., Bocherens, H., Witt, K.E., Samaniego Castruita, J.A., Velasco, M.S., et al. (2020). Ancient DNA suggests modern wolves trace their origin to a Late Pleistocene expansion from Beringia. Mol. Ecol. 29, 1596–1610.

van de Loosdrecht, M., Bouzouggar, A., Humphrey, L., Posth, C., Barton, N., Aximu-Petri, A., Nickel, B., Nagel, S., Talbi, E.H., El Hajraoui, M.A., et al. (2018). Pleistocene North African genomes link Near Eastern and sub-Saharan African human populations. Science 360, 548.

Maricic, T., Whitten, M., and Pääbo, S. (2010). Multiplexed DNA sequence capture of mitochondrial genomes using PCR products. PLoS One 5, e14004.

Martchenko, D., Chikhi, R., and Shafer, A.B.A. (2020). Genome Assembly and Analysis of the North American Mountain Goat (Oreamnos americanus) Reveals Species-Level Responses to Extreme Environments. G3 10, 437–442.

Martin, M. (2011). Cutadapt removes adapter sequences from high-throughput sequencing reads. EMBnet.journal 17, 10–12.

Mathieson, I., Lazaridis, I., Rohland, N., Mallick, S., Patterson, N., Roodenberg, S.A., Harney, E., Stewardson, K., Fernandes, D., Novak, M., et al. (2015). Genome-wide patterns of selection in 230 ancient Eurasians. Nature 528, 499–503.

McKenna, A., Hanna, M., Banks, E., Sivachenko, A., Cibulskis, K., Kernytsky, A., Garimella, K., Altshuler, D., Gabriel, S., Daly, M., et al. (2010). The Genome Analysis Toolkit: a MapReduce framework for analyzing next-generation DNA sequencing data. Genome Res. 20, 1297–1303.

Meyer, M., and Kircher, M. (2010). Illumina sequencing library preparation for highly multiplexed target capture and sequencing. Cold Spring Harb. Protoc. 2010, db.prot5448.

Narasimhan, V.M., Patterson, N., Moorjani, P., Rohland, N., Bernardos, R., Mallick, S., Lazaridis, I., Nakatsuka, N., Olalde, I., Lipson, M., et al. (2019). The formation of human populations in South and Central Asia. Science 365.

Okonechnikov, K., Conesa, A., and García-Alcalde, F. (2016). Qualimap 2: advanced multi-sample quality control for high-throughput sequencing data. Bioinformatics 32, 292–294.

Olalde, I., Allentoft, M.E., Sánchez-Quinto, F., Santpere, G., Chiang, C.W.K., DeGiorgio, M., Prado-Martinez, J., Rodríguez, J.A., Rasmussen, S., Quilez, J., et al. (2014). Derived immune and ancestral pigmentation alleles in a 7,000-year-old Mesolithic European. Nature 507, 225–228.

Patterson, N., Price, A.L., and Reich, D. (2006). Population Structure and Eigenanalysis. PLoS Genet. 2, e190.

Patterson, N., Moorjani, P., Luo, Y., Mallick, S., Rohland, N., Zhan, Y., Genschoreck, T., Webster, T., and Reich, D. (2012). Ancient admixture in human history. Genetics 192, 1065–1093.

Pedersen, M.W., Overballe-Petersen, S., Ermini, L., Sarkissian, C.D., Haile, J., Hellstrom, M., Spens, J., Thomsen, P.F., Bohmann, K., Cappellini, E., et al. (2015). Ancient and modern environmental DNA. Philos. Trans. R. Soc. Lond. B Biol. Sci. 370, 20130383.

Pedersen, M.W., Ruter, A., Schweger, C., Friebe, H., Staff, R.A., Kjeldsen, K.K., Mendoza, M.L.Z., Beaudoin, A.B., Zutter, C., Larsen, N.K., et al. (2016). Postglacial viability and colonization in North America’s ice-free corridor. Nature 537, 45–49.

Pinhasi, R., Meshveliani, T., Matskevich, Z., Bar-Oz, G., Weissbrod, L., Miller, C.E., Wilkinson, K., Lordkipanidze, D., Jakeli, N., Kvavadze, E., et al. (2014). Satsurblia: new insights of human response and survival across the Last Glacial Maximum in the southern Caucasus. PLoS One 9, e111271.

Plassais, J., Kim, J., Davis, B.W., Karyadi, D.M., Hogan, A.N., Harris, A.C., Decker, B., Parker, H.G., and Ostrander, E.A. (2019). Whole genome sequencing of canids reveals genomic regions under selection and variants influencing morphology. Nat. Commun. 10, 1489.

Posth, C., Renaud, G., Mittnik, A., Drucker, D.G., Rougier, H., Cupillard, C., Valentin, F., Thevenet, C., Furtwängler, A., Wißing, C., et al. (2016). Pleistocene Mitochondrial Genomes Suggest a Single Major Dispersal of Non-Africans and a Late Glacial Population Turnover in Europe. Curr. Biol. 26, 827–833.

Purcell, S., Neale, B., Todd-Brown, K., Thomas, L., Ferreira, M.A.R., Bender, D., Maller, J., Sklar, P., de Bakker, P.I.W., Daly, M.J., et al. (2007). PLINK: a tool set for whole-genome association and population-based linkage analyses. Am. J. Hum. Genet. 81, 559–575.

Racimo, F., Sikora, M., Vander Linden, M., Schroeder, H., and Lalueza-Fox, C. (2020). Beyond broad strokes: sociocultural insights from the study of ancient genomes. Nat. Rev. Genet.

Raghavan, M., Skoglund, P., Graf, K.E., Metspalu, M., Albrechtsen, A., Moltke, I., Rasmussen, S., Stafford, T.W., Jr, Orlando, L., Metspalu, E., et al. (2014). Upper Palaeolithic Siberian genome reveals dual ancestry of Native Americans. Nature 505, 87–91.

Renaud, G. (2018). libbam (Github).

Renaud, G., Slon, V., Duggan, A.T., and Kelso, J. (2015). Schmutzi: estimation of contamination and endogenous mitochondrial consensus calling for ancient DNA. Genome Biol. 16, 224.

Sánchez-Quinto, F., Schroeder, H., Ramirez, O., Avila-Arcos, M.C., Pybus, M., Olalde, I., Velazquez, A.M.V., Marcos, M.E.P., Encinas, J.M.V., Bertranpetit, J., et al. (2012). Genomic affinities of two 7,000-year-old Iberian hunter-gatherers. Curr. Biol. 22, 1494–1499.

Schiffels, S., Haak, W., Paajanen, P., Llamas, B., Popescu, E., Loe, L., Clarke, R., Lyons, A., Mortimer, R., Sayer, D., et al. (2016). Iron Age and Anglo-Saxon genomes from East England reveal British migration history. Nat. Commun. 7, 10408.

Sikora, M., Seguin-Orlando, A., Sousa, V.C., Albrechtsen, A., Korneliussen, T., Ko, A., Rasmussen, S., Dupanloup, I., Nigst, P.R., Bosch, M.D., et al. (2017). Ancient genomes show social and reproductive behavior of early Upper Paleolithic foragers. Science 358, 659–662.

Simpson, J.T., and Durbin, R. (2010). Efficient construction of an assembly string graph using the FM-index. Bioinformatics 26, i367–i373.

Sinding, M.-H.S., Gopalakrishan, S., Vieira, F.G., Samaniego Castruita, J.A., Raundrup, K., Heide Jørgensen, M.P., Meldgaard, M., Petersen, B., Sicheritz-Ponten, T., Mikkelsen, J.B., et al. (2018). Population genomics of grey wolves and wolf-like canids in North America. PLoS Genet. 14, e1007745.

Sinding, M.-H.S., Gopalakrishnan, S., Ramos-Madrigal, J., de Manuel, M., Pitulko, V.V., Kuderna, L., Feuerborn, T.R., Frantz, L.A.F., Vieira, F.G., Niemann, J., et al. (2020). Arctic-adapted dogs emerged at the Pleistocene–Holocene transition. Science 368, 1495–1499.

Skoglund, P., Storå, J., Götherström, A., and Jakobsson, M. (2013). Accurate sex identification of ancient human remains using DNA shotgun sequencing. J. Archaeol. Sci. 40, 4477–4482.

Skoglund, P., Northoff, B.H., and Shunkov, M.V. (2014). Separating endogenous ancient DNA from modern day contamination in a Siberian Neandertal. Proceedings of the.

Skoglund, P., Ersmark, E., Palkopoulou, E., and Dalén, L. (2015). Ancient wolf genome reveals an early divergence of domestic dog ancestors and admixture into high-latitude breeds. Curr. Biol. 25, 1515–1519.

Slon, V., Hopfe, C., Weiß, C.L., Mafessoni, F., de la Rasilla, M., Lalueza-Fox, C., Rosas, A., Soressi, M., Knul, M.V., Miller, R., et al. (2017). Neandertal and Denisovan DNA from Pleistocene sediments. Science 356, 605–608.

Søe, M.J., Nejsum, P., Seersholm, F.V., Fredensborg, B.L., Habraken, R., Haase, K., Hald, M.M., Simonsen, R., Højlund, F., Blanke, L., et al. (2018). Ancient DNA from latrines in Northern Europe and the Middle East (500 BC-1700 AD) reveals past parasites and diet. PLoS One 13, e0195481.

Suchard, M.A., Lemey, P., Baele, G., Ayres, D.L., Drummond, A.J., and Rambaut, A. (2018). Bayesian phylogenetic and phylodynamic data integration using BEAST 1.10. Virus Evol 4, vey016.

Thalmann, O., Shapiro, B., Cui, P., Schuenemann, V.J., Sawyer, S.K., Greenfield, D.L., Germonpré, M.B., Sablin, M.V., López-Giráldez, F., Domingo-Roura, X., et al. (2013). Complete mitochondrial genomes of ancient canids suggest a European origin of domestic dogs. Science 342, 871–874.

Upadhyay, M., Hauser, A., Kunz, E., Krebs, S., Blum, H., Dotsev, A., Okhlopkov, I., Bagirov, V., Brem, G., Zinovieva, N., et al. (2020). The First Draft Genome Assembly of Snow Sheep (Ovis nivicola). Genome Biol. Evol. 12, 1330–1336.

Vai, S., Sarno, S., Lari, M., Luiselli, D., Manzi, G., Gallinaro, M., Mataich, S., Hübner, A., Modi, A., Pilli, E., et al. (2019). Ancestral mitochondrial N lineage from the Neolithic “green” Sahara. Sci. Rep. 9, 3530.

Verdugo, M.P., Mullin, V.E., Scheu, A., Mattiangeli, V., Daly, K.G., Maisano Delser, P., Hare, A.J., Burger, J., Collins, M.J., Kehati, R., et al. (2019). Ancient cattle genomics, origins, and rapid turnover in the Fertile Crescent. Science 365, 173–176.

Wecek, K., Hartmann, S., Paijmans, J.L.A., Taron, U., Xenikoudakis, G., Cahill, J.A., Heintzman, P.D., Shapiro, B., Baryshnikov, G., Bunevich, A.N., et al. (2016). Complex Admixture Preceded and Followed the Extinction of Wisent in the Wild. Mol. Biol. Evol. 34, 598–612.

Weissensteiner, H., Pacher, D., Kloss-Brandstätter, A., Forer, L., Specht, G., Bandelt, H.-J., Kronenberg, F., Salas, A., and Schönherr, S. (2016). HaploGrep 2: mitochondrial haplogroup classification in the era of high-throughput sequencing. Nucleic Acids Res. 44, W58–W63.

Willerslev, E., Hansen, A.J., Binladen, J., Brand, T.B., Gilbert, M.T.P., Shapiro, B., Bunce, M., Wiuf, C., Gilichinsky, D.A., and Cooper, A. (2003). Diverse Plant and Animal Genetic Records from Holocene and Pleistocene Sediments. Science 1084114.

Willerslev, E., Davison, J., Moora, M., Zobel, M., Coissac, E., Edwards, M.E., Lorenzen, E.D., Vestergård, M., Gussarova, G., Haile, J., et al. (2014). Fifty thousand years of Arctic vegetation and megafaunal diet. Nature 506, 47–51.

Wu, D.-D., Ding, X.-D., Wang, S., Wójcik, J.M., Zhang, Y., Tokarska, M., Li, Y., Wang, M.-S., Faruque, O., Nielsen, R., et al. (2018). Pervasive introgression facilitated domestication and adaptation in the Bos species complex. Nat Ecol Evol 2, 1139–1145.

Yang, M.A., Gao, X., Theunert, C., Tong, H., Aximu-Petri, A., Nickel, B., Slatkin, M., Meyer, M., Pääbo, S., Kelso, J., et al. (2017a). 40,000-Year-Old Individual from Asia Provides Insight into Early Population Structure in Eurasia. Curr. Biol. 27, 3202–3208.e9.

Yang, Y., Wang, Y., Zhao, Y., Zhang, X., Li, R., Chen, L., Zhang, G., Jiang, Y., Qiu, Q., Wang, W., et al. (2017b). Draft genome of the Marco Polo Sheep (Ovis ammon polii). Gigascience 6, 1–7.

Yu, H., Spyrou, M.A., Karapetian, M., Shnaider, S., Radzevičiūte, R., Nägele, K., Neumann, G.U., Penske, S., Zech, J., Lucas, M., et al. (2020). Paleolithic to Bronze Age Siberians Reveal Connections with First Americans and across Eurasia. Cell.

Zhang, D., Xia, H., Chen, F., Li, B., Slon, V., Cheng, T., Yang, R., Jacobs, Z., Dai, Q., Massilani, D., et al. (2020). Denisovan DNA in Late Pleistocene sediments from Baishiya Karst Cave on the Tibetan Plateau. Science 370, 584–587.

Zheng, Z., Wang, X., Li, M., Li, Y., Yang, Z., Wang, X., Pan, X., Gong, M., Zhang, Y., Guo, Y., et al. (2020). The origin of domestication genes in goats. Science Advances 6, eaaz5216. Picard-tools.

