## Supplementary material for "Genome-scale sequencing and analysis of human, wolf and bison DNA from 25,000 year-old sediment": Sup-figures

**Figures S1 to S7**

**Tables S3 to S7**

**Figure S1. Sample descriptives:** A) Metagenomic prediction with centrifuge, B), Length distribution of the four aligned mammalian species C) Deamination pattern of the filtered reads of the four mapped genomes: *Ovis aries, Homo sapiens, Bos taurus* and *Canis lupus.*

**Figure S2. ADMIXTURE plots.** ADMIXTURE plots using K from 1 to 15 of the human dataset.

**Figure S3. $f_3$-Outgroup analysis. A)** Values of the $f_3$-outgroup statistics in the combination $f_3$(SAT29,X;Mbuti). Only combinations with more than 2,000 SNPs were included, green dots represent post-LGM samples, violet dots represent pre-LGM samples and red dots represent farmer samples. Error bars denote standard errors. **B**) Results of the $f_3$ (SAT29,Modern-population;Mbuti) combination.

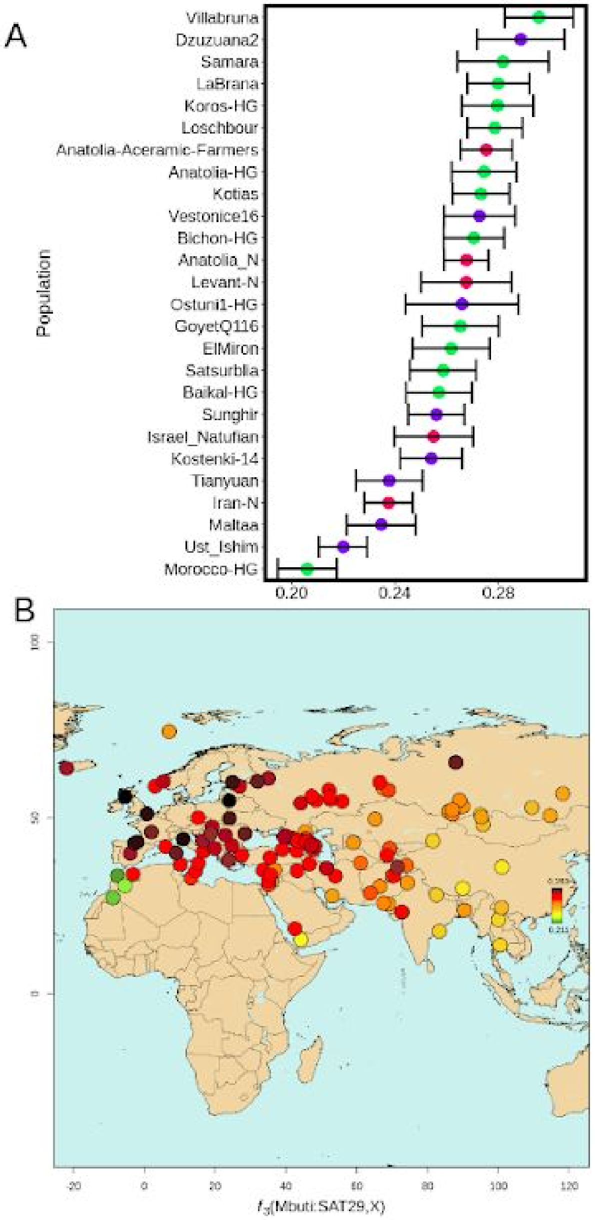

**Figure S4. The SAT29 *Canis* genome is symmetrically related to modern wolves and dogs, and the latter share drift to the exclusion of SAT29**. $f_4$-statistics evaluating the basal ancestry status of the SAT29 canid genome in relation to a set of 15 diverse Eurasian present-day wolves and dogs. Error bars denote ±3 standard errors. A) Statistics of the form $f_4$(CoyoteCalifornia,SAT29.*Canis*;A,B) are consistent with being distributed around 0, indicating that SAT29 is symmetrically related to present-day individuals. A few statistics fall close to or slightly outside $|Z| > 3$ (largest $|Z| = 3.43$), and involve wolves from Southwestern and South Asia (WolfSyria, WolfSaudiArabia, Wolf07Israel, Wolf19India). However, the values of these statistics are in the direction of SAT29 being closer to other wolves than to these wolves, and thus likely reflects some basal ancestry in these present-day wolves, e.g. due to admixture from non-wolf canids. B) Statistics of the form $f_4$(CoyoteCalifornia,A;B,SAT29.*Canis*) are all negative, indicating that all of the present-day wolves and dogs share genetic drift that is not shared with the SAT29 sample.

**Fig. S5. Principal component analysis on the *Ovis* and *Capra* dataset.** The analysis was performed on a distance matrix comparing per-sample profiles of all possible $f_4$-statistics of the form $f_4$(X,A;B,C) where X is the focal sample.. S12: PC 1-2 plot, S13: PC 3-4 plot, S14: PC 5-6 plot, S15: PC 7-8 plot, S16: PC 9-10 plot.

**Figure S6. Two microphotographs (A, B) from block sample SAT 15 14**. The microphotographs were taken adjacent to SAT 16 LS29, in cross- (XPL) and plane-polarized light (PPL), showing dominant components and processes. Typical anthropogenic components are bone (B), burnt bone (BB), charcoal (Ch) and flint (Fl). Note also the occurrence of rounded soil aggregates (yellow circles) that were transported into the cave from soils forming outside and that exhibit cross-striations of the clay component resulting from repeated wetting and drying cycles during their formation. Similarly, clay coatings in voids (blue arrow) result from water percolating through the sediment.

**Figure S7. Evidence for polymorphism in the SAT29 wolf mitochondrial sequence.** Sites that differ between the pre-LGM Armenian wolf TU10 and the major clade of wolf mitochondria were identified, and the read counts of the SAT29 sample at these sites displayed. Dark blue indicates SAT29 reads matching the major clade allele, while light blue indicates SAT29 matching the TU10 allele. Variants are stratified by mutational direction, with sites where derived mutations in SAT29 and TU10 can not be explained by C->T deamination on the left (black text), and sites where they could be on the right (gray text). A) Results for all reads. B) Results restricted to reads displaying evidence of deamination. These results suggest that the retrieved DNA derives from more than one wolf individual.

**Table S3. Human Mitochondria derived positions:** Derived positions of the SAT29 mitochondrial genome respect the rCRS sequence. Bold positions are the ones that have been accepted because the ratio derived/no derived was higher than ⅔.

| Position | Coverage | Alleles | Haplogroup |
|---|---|---|---|
| 73 | 7 | **0 A/7 G** | N |
| 263 | 24 | **2 A / 22 G** | N |
| 750 | 22 | **A 0 / G 22** | N |
| 1438 | 9 | **A 0 / G 9** | N |
| 2706 | 8 | **A 1 / G 7** | N |
| 4769 | 12 | **A 1 / G 11** | N |
| 7028 | 15 | **1 C / 14 T** | N |
| 8860 | 12 | **1 A / 11 G** | N |
| 11719 | 14 | **0 G / 14 A** | N |
| 12705 | 12 | **2 C / 10 T** | N |
| 14766 | 20 | **0 C / 20 T** | N |
| 15326 | 17 | **0 A / 17 G** | N |
| 16223 | 16 | 7 C / 9 T | N |
| 152 | 5 | 2 T / 3 C | private |
| 827 | 13 | 7 A / 6 G | private |
| 8655 | 14 | **4 C / 10 T** | private |
| 12451 | 15 | 6 A / 9 G | private |
| 12662 | 19 | **A 6 / 13 G** | private |
| 14218 | 17 | **5 T / 12 C** | private |
| 16189 | 14 | 7T/7C | private |
| 16519 | 8 | **0 T / 8 C** | hotspot |

**Table S6. $f_4$-outgroup statistics results of SAT29 with human ancient individuals:** $f_4$-statistics results in the combination $f_4$(Dzudzuana2,X;SAT29,Mbuti).

| Pop A | Pop B | Pop C | Pop D | $f_4$-value | SD | Z score | SNPs |
|---|---|---|---|---|---|---|---|
| Dzuzuana2 | Anatolia_N | SAT29 | Mbuti | 0.007312 | 0.003279 | 2.230 | 4463 |
| Dzuzuana2 | Tianyuan | SAT29 | Mbuti | 0.010573 | 0.004824 | 2.192 | 3673 |
| Dzuzuana2 | Vestonice16 | SAT29 | Mbuti | 0.009875 | 0.005098 | 1.937 | 3118 |
| Dzuzuana2 | Kotias | SAT29 | Mbuti | 0.007575 | 0.004297 | 1.763 | 4483 |
| Dzuzuana2 | Satsurblia | SAT29 | Mbuti | 0.005941 | 0.005189 | 1.145 | 3198 |
| Dzuzuana2 | GoyetQ116 | SAT29 | Mbuti | 0.010481 | 0.005157 | 2.032 | 3240 |
| Dzuzuana2 | Koros-HG | SAT29 | Mbuti | -0.000125 | 0.004673 | -0.027 | 3327 |
| Dzuzuana2 | ElMiron | SAT29 | Mbuti | -0.002115 | 0.005708 | -0.371 | 2659 |
| Dzuzuana2 | LaBrana | SAT29 | Mbuti | 0.003839 | 0.004445 | 0.864 | 4222 |
| Dzuzuana2 | Iran-N | SAT29 | Mbuti | 0.010591 | 0.003743 | 2.830 | 4285 |
| Dzuzuana2 | Villabruna | SAT29 | Mbuti | -0.004202 | 0.004553 | -0.29 | 1954 |
| Dzuzuana2 | Ostuni1-HG | SAT29 | Mbuti | -0.001887 | 0.007509 | -0.251 | 1369 |
| Dzuzuana2 | Levant-N | SAT29 | Mbuti | 0.003945 | 0.013064 | 0.30 | 507 |
| Dzuzuana2 | Loschbour | SAT29 | Mbuti | 0.003366 | 0.003852 | 0.874 | 4438 |
| Dzuzuana2 | Morocco-HG | SAT29 | Mbuti | 0.019854 | 0.004474 | 4.438 | 4021 |
| Dzuzuana2 | Samara | SAT29 | Mbuti | 0.005471 | 0.006641 | 0.824 | 2003 |
| Dzuzuana2 | Kostenki-14 | SAT29 | Mbuti | 0.004512 | 0.004462 | 1.011 | 4257 |
| Dzuzuana2 | Maltaa | SAT29 | Mbuti | 0.011467 | 0.005187 | 2.211 | 4257 |
| Dzuzuana2 | Sunghir | SAT29 | Mbuti | 0.009413 | 0.004071 | 2.312 | 4484 |
| Dzuzuana2 | Ust_Ishim | SAT29 | Mbuti | 0.015006 | 0.003692 | 4.065 | 4476 |
| Dzuzuana2 | Bichon-HG | SAT29 | Mbuti | 0.004794 | 0.004296 | 1.116 | 4476 |
| Dzuzuana2 | Baikal-HG | SAT29 | Mbuti | 0.002936 | 0.004688 | 0.626 | 3562 |
| Dzuzuana2 | Anatolia-Acerami c-Farmers | SAT29 | Mbuti | 0.005734 | 0.003898 | 1.471 | 4272 |
| Dzuzuana2 | Anatolia-HG | SAT29 | Mbuti | 0.007690 | 0.004857 | 1.583 | 3484 |
| Dzuzuana2 | Israel_Natufian | SAT29 | Mbuti | 0.004809 | 0.005557 | 0.865 | 2411 |

**Table S7. Human mitochondrial mapping differences:** Mitochondrial mapping differences between the three different bins: A) All reads, B) Deaminated reads, C) Non deaminated reads. The discordant positions are shown in this table

| Positions | All Reads | Non deaminated reads | Deaminated reads |
|---|---|---|---|
| 497 | 7C/6T | 1T/3C | 2T/1C |
| 3480 | 5G/5A | 5G/1A | 3A |
| 11914 | 11G/3A | 2G/3A | 7G |
| 14905 | 9G/8A | 4G/7A | 4G |
